# S-Alkyl-Phosphorothioate Modifications Reduce Thermal and Structural Stability of DNA Duplexes

**DOI:** 10.1101/2025.09.30.679560

**Authors:** Soumya Chandrasekhar, Rachel Bricker, Fatemeh Fadaei, Thorsten-Lars Schmidt

**Author notes:** Contributed equally (Soumya Chandrasekhar, Rachel Bricker).

## Abstract

While phosphorothioate (PS) oligonucleotides are usually used in therapeutic applications, they also offer the cheapest and synthetically most straightforward route to introduce hydrophobic modifications for applications in structural DNA nanotechnology and biophysics. For this, the sulfur atom is S-alkylated with alkyl iodides, enabling a hydrophobically tunable interface of DNA nanostructures with lipid bilayers. While longer and more alkyls per helical turn should lead to stronger interactions with lipid membranes, we found that excessive S-alkylations strongly inhibit hybridization of oligonucleotides to their complementary strands and decrease their melting temperature, despite a reduction in electrostatic repulsion between the two strands. Moreover, both the type and placement of alkyl modifications influence the melting temperature. Atomistic molecular dynamics simulations reveal two complementary mechanisms that explain the experimental findings. First, S-alkylated oligonucleotides are more compact and less dynamic than unmodified ones, likely inhibiting their ability to hybridize to their complementary strands. Second, S-alkyls in double-stranded DNA promote defect formation due to alkyl modifications having hydrophobic interactions with other alkyl groups and nucleobases, therefore reducing the thermal and structural stability of alkylated DNA duplexes. This study serves as a practical guide for tuning hydrophobicity while maintaining structural stability in membrane-interfacing DNA nanostructures.

## Introduction

Chemical modifications of natural nucleic acids and artificial, synthetic nucleic acid analogues have been explored for decades, mostly for applications in biomedicine such as therapeutics that modulate gene expression.^[1]^ Modifications can be introduced at various positions including phosphate, sugar, or nucleobases. Synthetic backbone analogues or modifications include phosphorothioates (PS), methylphosphonates, peptide nucleic acids (PNA), Glycol nucleic acids (GNA), 2’-O-Methyl, 2’-Fluoro, morpholino, locked nucleic acids, spiegelmers, and many more.^[2]^ Such modified nucleic acids are usually degraded slower by nucleases and can show improved uptake or pharmacokinetics than standard oligonucleotides. Such pharmaceutical or biomedical applications are, however, not the scope of this paper and are reviewed elsewhere. ^[1,2]^

Another important application for both standard and modified nucleic acids is structural DNA nanotechnology.^[3]^ DNA is arguably the most programmable and addressable biomolecule, and enables the creation of complex, functionalizable nanostructures.^[4]^ Both the deletion of charges and DNA modifications with hydrophobic groups^[5]^ enables interactions between DNA and lipid membranes for: biosensing;^[6,7]^ artificial cell development;^[8]^ creating DNA nanopores that function as artificial ion channels;^[9–13]^ scaffolded DNA-lipid nanodiscs;^[14]^ understanding the mechanism of viral entry;^[15]^ inducing curving and deformation of lipid membranes;^[16,17]^ regulating membrane fusion reactions;^[18,19]^ controlling shapes of liposomes by acting as nanotemplates;^[20]^ or mimicking enveloped virus particles for biomedical applications.^[21]^

Common hydrophobic DNA modifications that insert themselves into the lipid bilayer include cholesterol,^[22,23]^ tocopherol,^[24]^ phospholipids,^[21,25]^ porphyrins,^[26–28]^ pyrene,^[29]^ and S-alkylated PS.^[10,14]^ From the various chemistries and positions on oligonucleotides that can be modified with hydrophobic groups, S-alkylation of PS oligonucleotides stands out as the most tunable, cheapest, and experimentally most straightforward option. While many modifications are commercially available, they can significantly increase the cost per oligonucleotide. In-house DNA synthesis requires elaborate, multi-step organic synthesis for the preparation of phosphoramidites and purchasing and maintaining DNA synthesizers. On the other hand, PS oligonucleotides are commercially available from all big oligonucleotides vendors and are the cheapest backbone modification, adding only ∼3 USD per modification for standard ∼100 nmol synthesis scales.

In PS oligonucleotides, a sulfur atom replaces one of the non-bridging phosphate oxygens. The sulfur atom is introduced during oligonucleotide synthesis with the sulfurizing Beaucage reagent in the oxidation step ^[30]^ and produces diastereomer mixtures. Stereopure oligonucleotides can be obtained,^[31]^ but only at much higher synthetic effort. Since sulfur is larger, more polarizable, and more nucleophilic than oxygen, the negative charge in a PS is located on the sulfur atom.^[32]^ Therefore, alkyl iodides can be used for a selective and quantitative S-alkylation ^[33]^ of the sulfur in a bimolecular nucleophilic substitution (Figure 1A), resulting in a charge-neutral and hydrophobic backbone segment.

**Figure 1.**
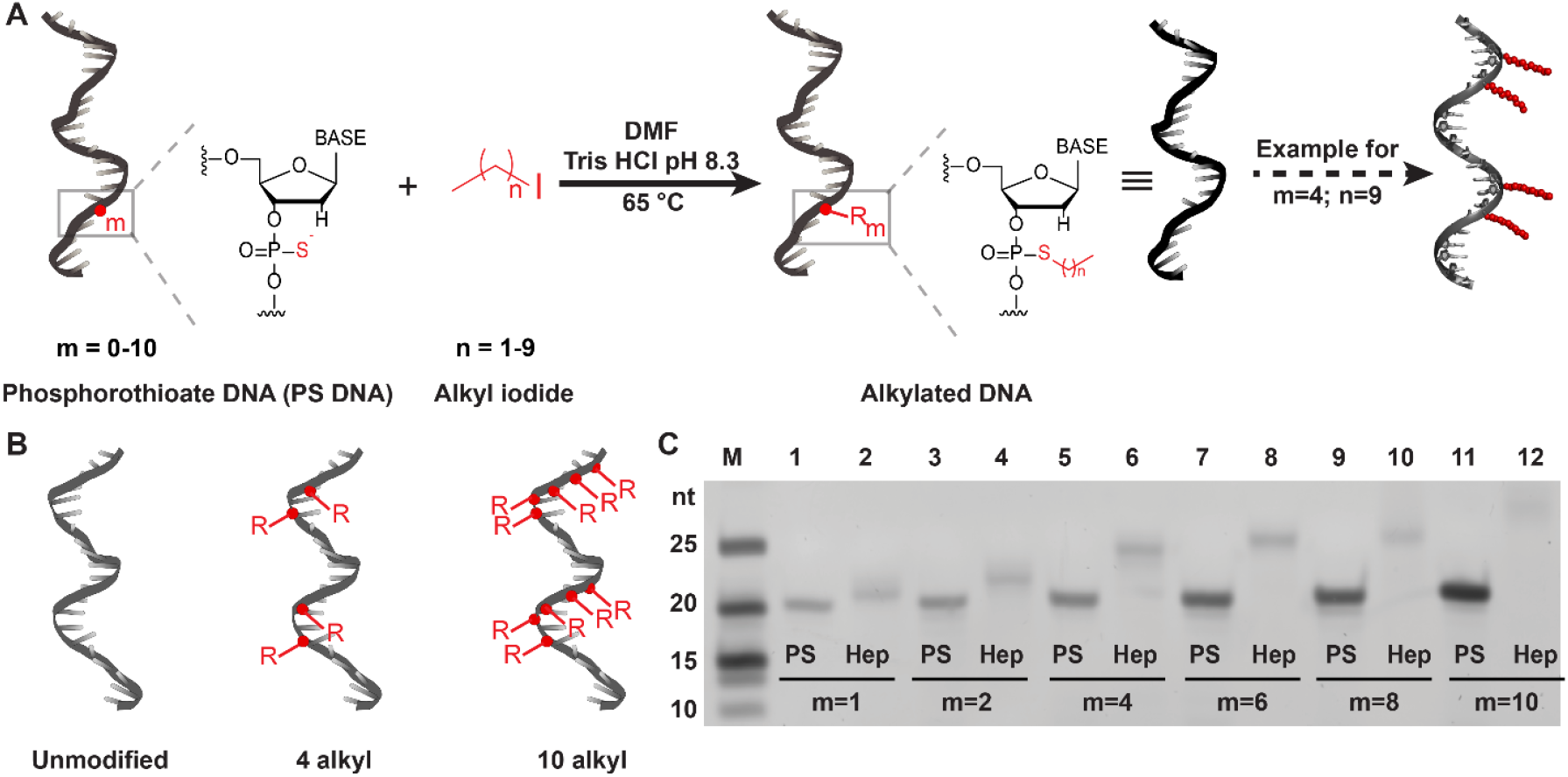
Synthesis and characterization of alkylated single-stranded DNA. (A) S-alkylation of PS DNA (grey strand), where “m” indicates the number of PS in the DNA backbone and “n” the number of methylenes in the alkyl chain. The black strand represents the alkylated product with values of m=0-10 and n=1-9. The model on the right shows an oligonucleotide with four decyl modifications (m=4, n=9) for illustrative purposes. (B) Illustrations of the modification designs: unmodified DNA, DNA with four alkyl modifications, and DNA with 10 alkyl modifications. Modifications were placed on one side of a duplex to create defined interfaces with lipids. (C) Denaturing PAGE of oligonucleotides with m=1-10 PS, before and after reaction with heptyl iodide (n=6).

As PS groups can replace any bridging phosphates, or even all, and can be modified with alkyls of different lengths, the orientation and hydrophobicity of these groups can be finely tuned with respect to the rest of a DNA nanostructure. This modification is therefore frequently used in DNA nanostructures including: synthetic DNA nanopores^[10,11]^ that insert themselves into a lipid membrane; DNA tetrahedra that interact with proteins^[34,35]^ or for the investigation of the immunological response of mammalian primary splenic cells to CpG sequences.^[36]^ In our lab, we synthesized a DNA-lipid nanodisc as a lipid bilayer mimetic for studying membrane proteins.^[14]^ For this, we created a double-stranded DNA minicircle in which we alkylated one to two PS groups with different alkyls ranging from ethyl to decyl that enveloped a lipid bilayer and found longer alkyls to hold the bilayer better.

Subsequent coarse-grained molecular dynamics simulations, on the other hand, suggested that charge neutralization of the DNA scaffold with many short alkyl groups will result in stronger interactions with lipids and more stable bilayers than nanodiscs with few longer S-Alkyls.^[37]^ In that report, the prediction was not experimentally tested, though.

In this study, we therefore systematically increased the number of PS to test if extreme S-alkylation is an experimentally viable strategy for DNA nanotechnology, and to potentially produce more stable DNA-lipid nanodiscs than in our initial design.^[14]^ Specifically, we investigated the effect of S-alkylation of PS oligonucleotides on DNA duplex stability by experimentally measuring their hybridization propensity and melting temperatures (*T*_*m*_). Although we originally expected that a charge neutralization on the backbone might lead to stronger DNA binding due to a reduced electrostatic repulsion, we found the opposite effect. Atomistic molecular dynamics (MD) simulations suggest mechanisms that explain the detrimental effect on the structure of single- and double-stranded DNA with S-alkyl PS modifications.

## Main Text

### S-alkylation of oligonucleotides

We alkylated a 21-base oligonucleotide without secondary structures and varied the number of PS modifications (m) between 1-10. Thus, 5-50% of the backbone charges were neutralized. Alkyl chain lengths (n=number of methylene units) were varied from ethyl (n=1) to decyl (n=9). Commercial PS oligonucleotides were subjected to S_N_2 reactions with alkyl iodide by a method adapted from Ref. ^[38]^ (Figure 1A). Note that after solvent evaporation, the pellet could only be fully solubilized in a hot EDTA (disodium ethylenediaminetetraacetic acid) solution.^[39]^ To interface with bilayers,^[14]^ all alkyls were designed to be on the same side of the DNA helix (Figure 1B).

Figure 1C shows a denaturing polyacrylamide gel electrophoresis (PAGE) of the PS strands before and after alkylation. Alkylation reduces electrophoretic mobility, which can be attributed to the increased mass and decreased negative charge. Note that all lanes in Figure 1C contain the same amount of photometrically quantified DNA, but appear at different intensities, which we attribute to different staining efficiency with the fluorescent SYBR dye. Additionally, we characterized the reaction products by reverse phase-high performance liquid chromatography (RP-HPLC), also confirming successful product formation (Figure S1, Supporting Information).

Three trends can be observed in Figure 1C. First, a higher number of PS groups increase band intensity. While the number of charges and potential ionic interactions with the cationic dye remains the same, PS is a softer Lewis base that might have a higher binding affinity with the dye than standard phosphates. Second, alkylation generally weakens band intensities, likely due to reduced electrostatic interaction with the stain. Third, increasing the length of alkyl chains from ethyl to decyl also improves staining, likely due to stronger hydrophobic interactions with the dye or an increase in its fluorescence intensity in a less polar environment. These effects make quantification of band intensities difficult, and gels can only be interpreted qualitatively.

### S-alkylation inhibits hybridization

We then hybridized these alkylated DNA with a complementary 147-nucleotide circular scaffold.^[14]^ For this, the single-stranded minicircle was annealed with seven identical complementary 21-mer alkylated oligonucleotides. The oligonucleotides were modified at either 2 or 10 positions with either butyl or heptyl chains (Figure 2A). We used the corresponding non-alkylated oligonucleotides as a control (Figure S2, Supporting Information) and characterized the products by native PAGE to analyze binding properties.

**Figure 2.**
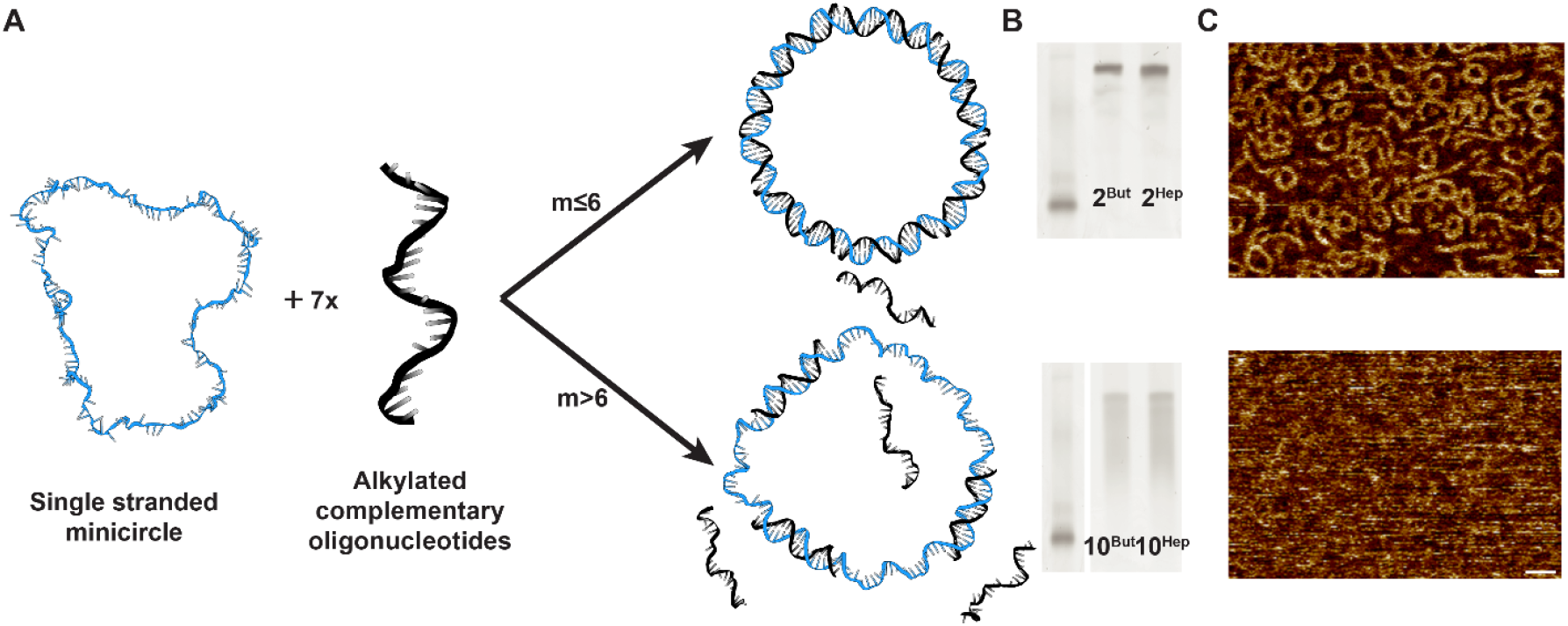
(A) Synthesis of double-stranded alkylated minicircles with varying extents of backbone modifications. (B) Native PAGE of double-stranded circles with 2 (top) or 10 (bottom) alkyls per oligonucleotide. (C) AFM images of intact double-stranded circles (two butyls per oligonucleotide) and partially denatured circles (10 butyls per oligonucleotide); scale bar = 20 nm.

As seen in Figure 2B-C, alkylated oligonucleotides form duplexes only when few alkyls are introduced. When 50% of the oligonucleotide backbone charges are neutralized with butyl or heptyl chains, we observed that the yield of the double-stranded rings was poor. Additionally, the entire lane appeared as a smear, indicating that hybridization was weakened. Contrary, we anticipated stronger hybridization due to decreased electrostatic repulsions, similar to trends seen in neutral peptide-nucleic acids.^[40]^ However, AFM images also show intact double-stranded rings only in cases where m≤6 per 21-mer (Figure 2C). Thus, we hypothesize that increasing S-alkylation decreases the *T*_*m*_ of the respective duplex.

### S-alkylation reduces *T*_*m*_

The *T*_*m*_ is defined as the temperature where 50 % of the base pairs in the DNA are denatured. To understand the effects of PS alkylation on the *T*_*m*_, we synthesized 21-base oligonucleotides alkylated with 1-10 groups: either butyl, pentyl, or heptyl. In denaturing PAGE gels, alkylated strands have a decreased electrophoretic mobility (Figure 1C, Figure S3, Supporting Information). Next, alkylated oligonucleotides were annealed with an unmodified complementary strand in the presence of 5 mM MgCl_2_ and 1X SYBR green I for the fluorescence detection of double-stranded DNA in a real-time PCR cycler. Controls included DNA with 1-10 PS modifications and non-PS DNA.

*T*_*m*_ measurements are shown in Figure 3. The measured *T*_*m*_ of the regular phosphate 21-base DNA duplex was 70 °C. We observed increasing the number of backbone PS slightly decreases the *T*_*m*_, consistent with a recent report.^[41]^ This can be attributed to the alteration in the electronic distribution and spatial configuration of the DNA backbone due to replacement of oxygen by sulfur.

**Figure 3.**
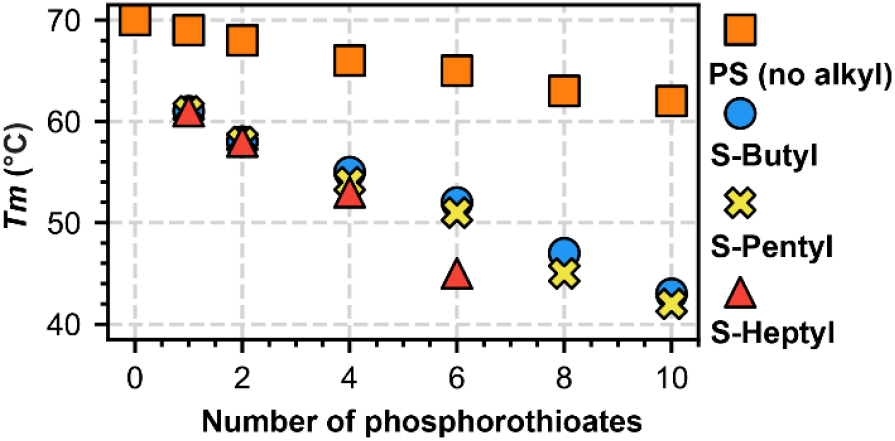
Melting temperatures before and after S-alkylation. Squares: *T*_*m*_ of strand with n PS before modification. Analysis with scatter plot indicating *T*_*m*_ of alkylated strand, varying from m=1-10 and n= 3, 4 and 6 (=butyl, pentyl, and heptyl).

When the backbone was neutralized, the *T*_*m*_ reduced much more, irrespective of the alkyl chain length. Increasing the number of backbone modifications reduced *T*_*m*_ to the point where the *T*_*m*_ could not be clearly determined anymore (raw data: Figure S4, Supporting Information). Different concentrations of SYBR green had no significant influence on the *T*_*m*_ of samples (Figure S5, Supporting Information).

PNA or morpholino oligonucleotides are also charge-neutral, but have a higher melting temperature than standard oligonucleotides, in part due to a reduced electrostatic repulsion. In S-alkylated PS oligonucleotides, however, the advantage of charge neutralization is apparently overcompensated by the detrimental hydrophobic interactions between alkyls and nucleobases that are absent in PNA and morpholino analogues. A similar *T*_*m*_ decrease is also observed in charge-neutral methylphosphonates.^[42]^

Next, the alkyl chain length influenced the *T*_*m*_ much less than their number. Differences became noticeable only if six or more modifications were introduced, and longer alkyls had a slightly decreased *T*_*m*_. We conclude that excessive charge neutralization and increasing alkyl chain length in a DNA backbone dramatically reduces its capability to hybridize with its complement.

### Relative position of alkyl groups

Next, we investigated the influence of the relative placement of the alkylations on the *T*_*m*_. Oligonucleotides with four PS were designed with the patterns (i, ii, iii) shown in Figure 4A and were modified with either butyl, heptyl, or decyl.

**Figure 4.**
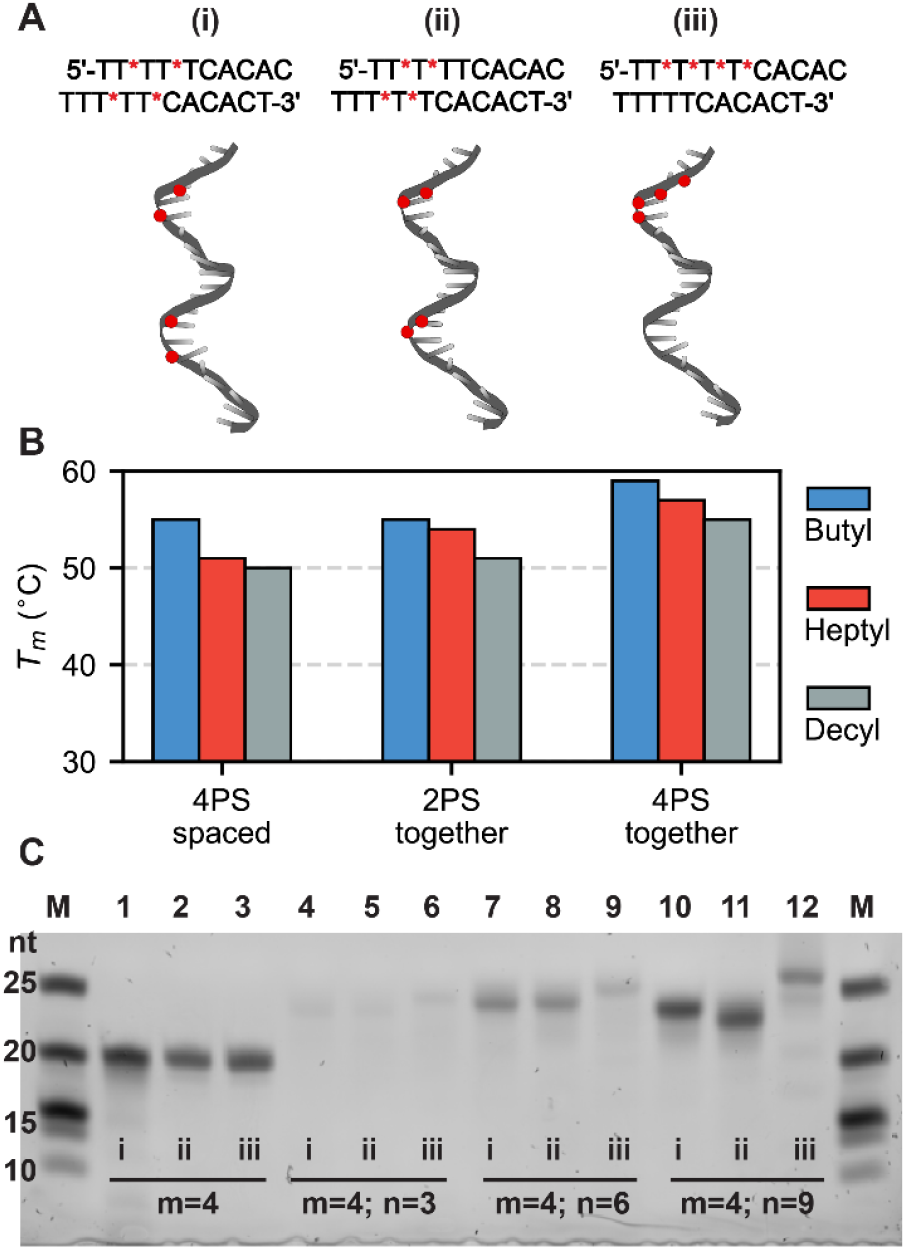
(A) Designs of 4 PS DNA with varying placements of PS groups. (B) Bar graph indicating *T*_*m*_ changes upon changing placement of the PS groups for different lengths of the alkyl chain. (C) Denaturing PAGE to show the modification of DNA (i), (ii), and (iii) with alkyl chains of different lengths.

Comparing their *T*_*m*_ (Figure 4B), the spaced patterns (i, ii) have a lower *T*_*m*_ than the variant with consecutive alkyl chains (iii). We hypothesize that backbone neutralization reduces *T*_*m*_ in a similar mechanism as a mismatched base. Indeed, mfold^[43]^ predictions of the *T*_*m*_ of duplexes with different mismatch patterns show the same trend: if mismatches are spaced apart, the *T*_*m*_ is lower than if mismatches are placed consecutively (Figure S6, Supporting Information). This suggests that both mismatches and alkylations have a destabilizing effect on the cooperative binding of neighboring base pairs.

The reduction of the electrophoretic mobility in denaturing PAGE (Figure 4C) indicates successful alkylation. Again, all lanes contain the same amount of DNA and band intensity differences are attributed to different staining efficiencies. Although each molecule has identical molecular weight and charge, the electrophoretic mobility is reduced when alkyl groups are placed together (iii) compared to spaced-out (i, ii). This is most clearly visibly in lanes 10-12 in Figure 4C. Without alkylation, the different PS placement patterns did not influence the *T*_*m*_ or electrophoretic mobility significantly (Figure S7, Supporting Information). This suggests that the spaced-out alkylations (i, ii) facilitate oligonucleotide compaction.

A reduction in the *T*_*m*_ means that the equilibrium between double- and single-stranded DNA is shifted towards the single strands. This can be caused by a destabilization of the double strand, a stabilization of the single strands, or both. We hypothesize that both effects play a role here, and that the compaction of alkylated single-stranded DNA is caused by hydrophobic interactions as indicated in the abstract figure. Thus, increasing the number of alkyls reduces their capability to hybridize to its complementary strand.

### MD simulations of single strands

To better understand the molecular origins of the experimental effects of alkylation, we performed atomistic MD simulations of unmodified (a) and alkylated (b-e) models of single-stranded oligonucleotides (Figure 5A). For models (b-d) we chose decyl groups since these had the greatest effects in *T*_*m*_ and gel experiments. Model (e) has ethyl groups to quantify the influence of the chain length. The models were simulated for 600 ns at constant ambient temperature using the CHARMM36 force field.

**Figure 5.**
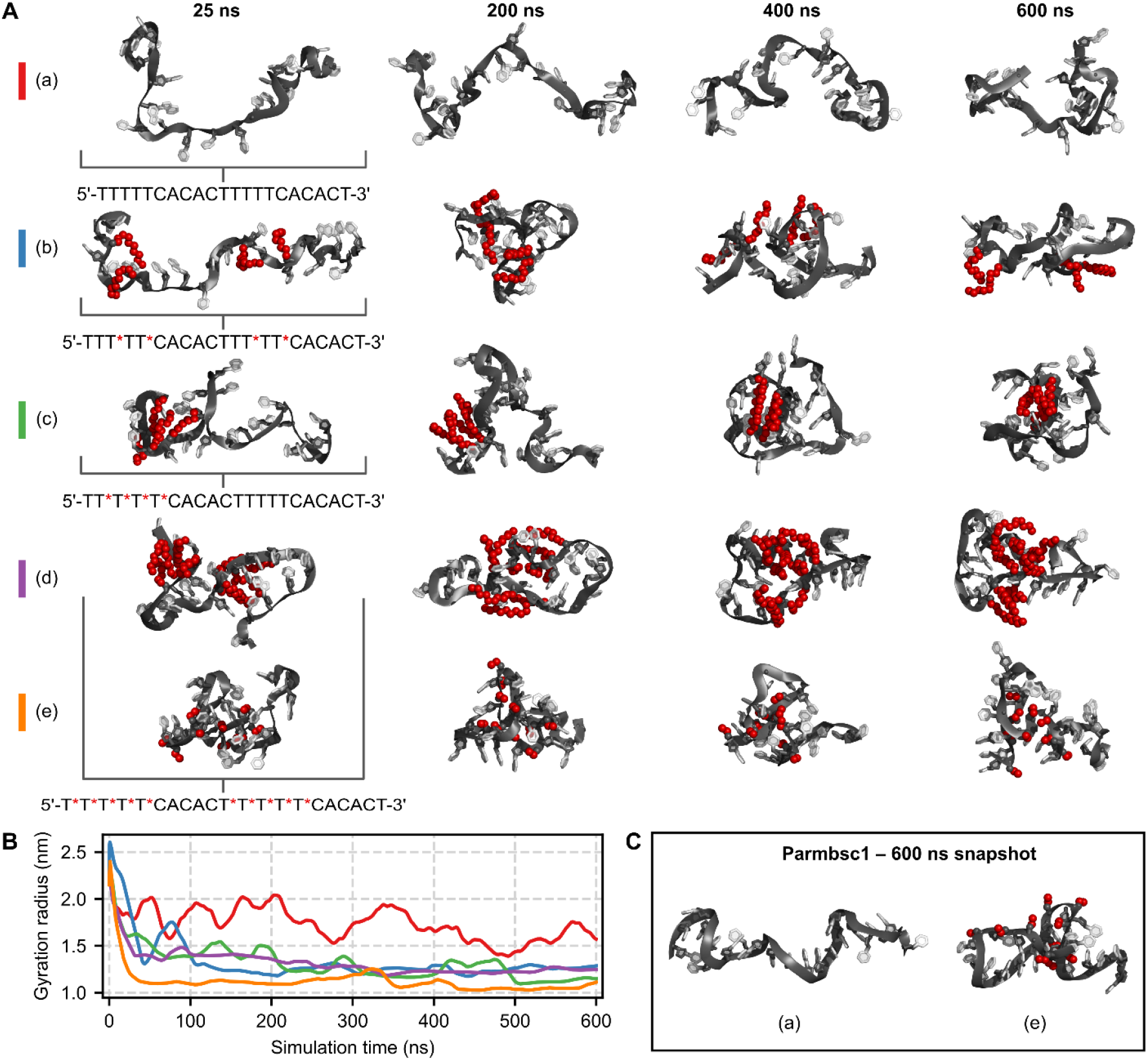
Atomistic MD simulations of unmodified and alkyl-modified single strands using either the CHARMM36 (A-B) or Parmbsc1 (C) force field. (A) Simulation snapshots at different times of the molecular dynamics trajectory. The methyl and methylene groups are shown as red spheres; red asterisks (*) in the sequences indicate alkylated bridging phosphates. (B) Radii of gyration of models (a-e) plotted as a function of simulation time; panel (A) show the color code. (C) Parmbsc1-based simulation snapshots at the 600 ns time step.

Snapshots of the models at various simulation times are shown in Figure 5A. To assess the compactness of each model, their gyration radius was calculated for each configuration of the simulation trajectory (Figure 5B). The average gyration radius values for models (a) through (e) are, respectively, 1.67 ± 0.23 nm, 1.25 ± 0.05 nm, 1.22 ± 0.10 nm, 1.25 ± 0.05 nm, and 1.10 ± 0.07 nm. The first 200 ns are excluded from the averages because the models are relaxing from their artificial initial structure: one strand of an idealized B-DNA helix.

The unmodified model (a) exhibits the largest average gyration radius value because the alkyl-modified models collapse to bury hydrophobic groups in their core, shielding them from water. Supplementary Movie 1 illustrates this. Additionally, the standard deviation in average gyration radius is largest for the unmodified model (a), indicating more dynamic conformational changes.

That is, the unmodified model tends to fluctuate between extended and compacted conformations, whereas the alkyl-modified models predominantly stay in a compact conformation.

The simulations indicate that the extent of compaction in alkyl-modified models is influenced by modification type. That is, the ethyl modified model (e) has the shortest average gyration radius. Intuitively, when the chains are short in length, more folding is required for the chains to be close enough to interact. If the first 200 ns are excluded, there is no clear correlation between the extent of compactness and either the placement or number of modifications.

In contrast, experiments indicate that the number of alkyl chains influences melting temperature more than their length, and that shorter chains lead to higher melting temperatures. Therefore, the short gyration radius of the oligonucleotides cannot be the only factor influencing the experimentally observed changes in *T*_*m*_, suggesting changes in the double-stranded DNA as well.

While force fields for DNA have become better in the last decades, no model is perfect. ^[44,45]^ To examine the influence of the force field, we repeated simulations with Parmbsc1, another popular force field for nucleic acids,^[46]^ Figure 5C shows final simulation snapshots of models (a) and (e). Parmbsc1 is known to over-estimate base stacking,^[47]^ causing unmodified single-stranded DNA to maintain an unrealistic, extended helical shape with little kinking and compaction. Nevertheless, the alkylated model (e) also compacts in Parmbsc1-based simulations, confirming that the alkyl groups facilitate folding in single-stranded DNA. Snapshots and gyration radii analyses of the Parmbsc1-based simulations are shown in Figure S8, Supporting Information.

### MD simulations of double strands

We next explored how alkylation influences double-stranded DNA. Previous MD simulations show a distortion of the DNA structure when the last six phosphates in both strands are alkylated.^[39]^ However, it is unclear how alkylation of only one strand, such as in DNA lipid nanodiscs ^[14,37]^ or DNA origami staples strand designs, impacts the DNA structure. Thus, we simulated the duplexes of strands (a) through (e) from Figure 5 at constant ambient temperature. The results are shown in Figure 6; data for models (c) and (e) are shown in Figure S9, Supporting Information. Once again, two different force fields were used: either Parmbsc1 (Figure 6A-C) or CHARMM36 (Figure 6D-F).

**Figure 6.**
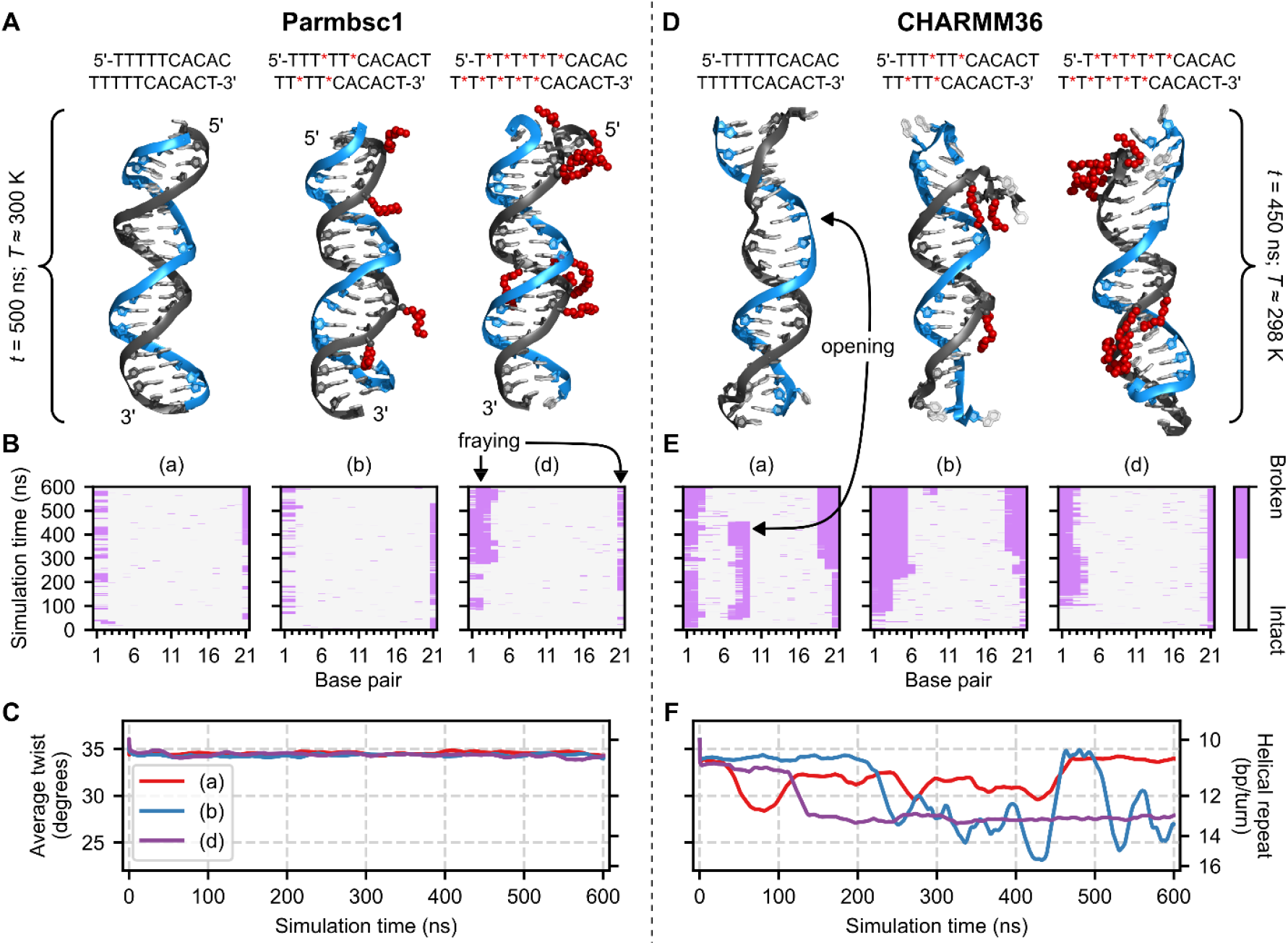
Constant temperature MD simulations of unmodified and alkyl-modified double-stranded DNA: the duplexe of strands (a), (b), and (d) from Figure 5. The left column (A-C) shows Parmbsc1-based simulation results, the right (D-F) CHARMM36-based. (A, D) Simulation snapshots at simulation time *t* and system temperature *T*. The models correspond to the plots in (B, E) below. (B, E) The 2D color plots show hydrogen bonding between Watson-Crick base pairs throughout the duplex for each frame. Light gray indicates hydrogen bonding, and light purple broken hydrogen bonding. Base pair 1 is the topmost base pair in the models shown in (A, D). (C, F) Time evolution of average twist, excluding the three terminal base pairs on each end of the duplex. The secondary *y*-axis shows the helical repeat, which was calculated from the average twist: (helical repeat) = 360°/(average twist).

Figure 6B, E shows color maps of intact or broken hydrogen bonds for each Watson-Crick base pair throughout the simulations. The CHARMM36-based simulation of model (d) exhibits backbone distortion while maintaining internal base pairing interactions (Figure 6D). Thus, hydrogen bonding analysis alone does not capture all distortions and another structural feature, twist, was analyzed. Figure 6C, F shows the time evolution of the average twist between neighboring base pairs, excluding the three terminal base pairs at each end of the duplex because of fraying.

The Parmbsc1-based simulations reveal that model (b) generally behaves like the unmodified model (a). In contrast, model (d) shows a higher degree of broken base pairs due to significant end-fraying. This distinction between alkyl-modified models (b) and (d) can be attributed to the greater count of alkyl groups, especially at the adenine–thymine rich terminal, in (d).

Unsurprisingly, all simulations show fraying of the ends ^[48]^ as no restraints were introduced in production simulations to prevent it. In addition, the CHARMM-based simulation of the unmodified model (a) shows unphysical internal base pair opening (Figure 6D-E and Supplementary Movie 2), consistent with a previous report.^[49]^ Therefore, the Parmbsc1 force field is considered the better model for the mechanical properties of double-stranded DNA.^[45]^

### Simulated melting of double strands

The duplexes remained mostly intact in the isothermal Parmbsc1-based simulations (Figure 6A-C), while experimentally these are not stable. This discrepancy can be explained by the low simulation temperature and the short time scales. Moreover, the initial structure is an idealized B-DNA helix, which might experimentally not form or only in small yields. Regardless, the simulations demonstrate a difference between model (d) and the unmodified model (a). To amplify the differences, we extended the simulations and linearly increased the system temperature from 300 K to 440 K within 1.2 μs using the “simulated annealing” option available in GROMACS (Figure 7).

**Figure 7.**
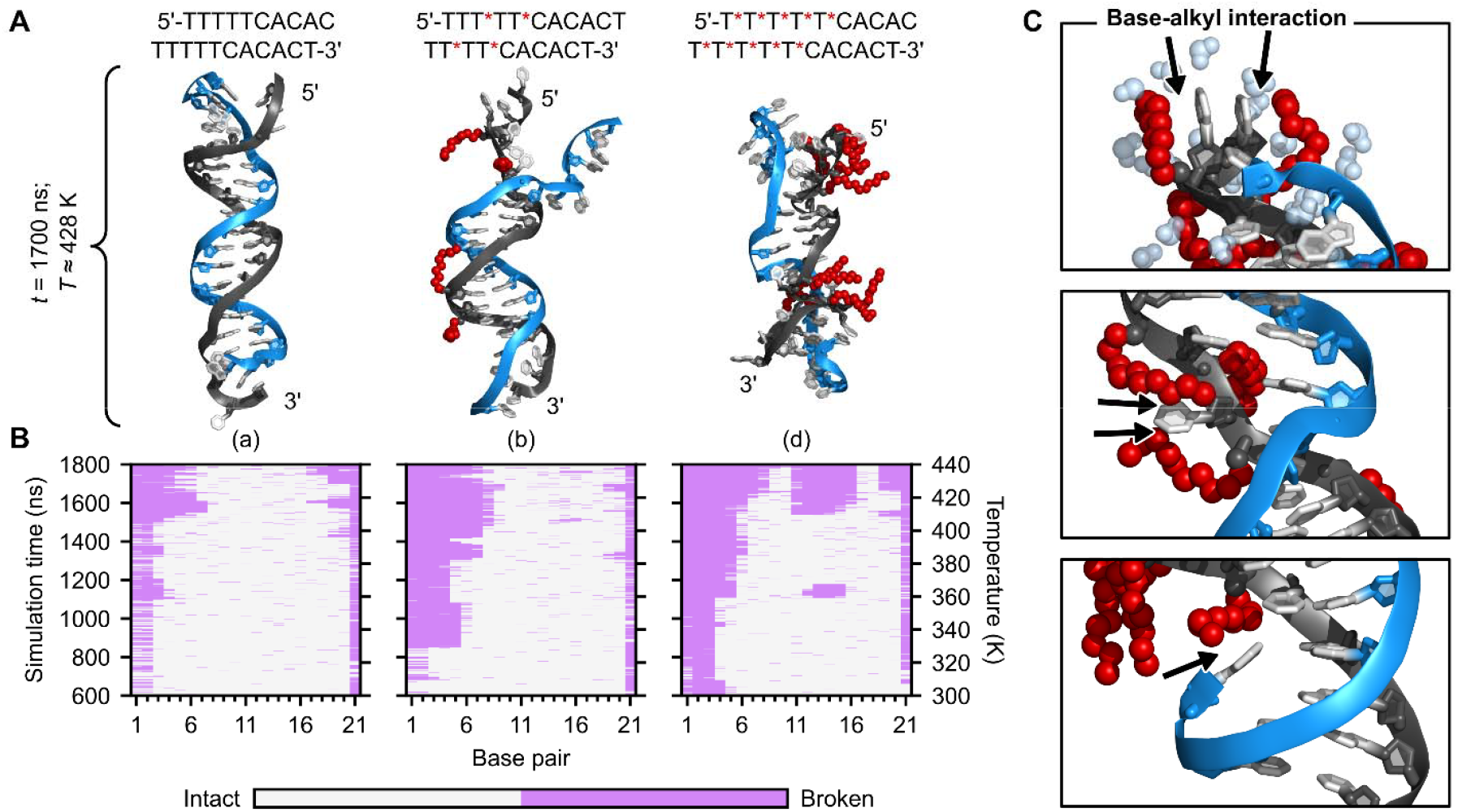
Effects of high temperatures. The Parmbsc1-based MD simulations shown in Figure 6A-C were extended with gradual heating of the system. (A) Simulation snapshots after 1700 ns at approximately 428 K. (B) The 2D color plots show intact (light grey) and broken (light purple) base pairs for each frame. Since the initial configurations of the heating simulations are the last configurations of the constant temperature ones, the starting time is marked as 600 ns. The secondary *y*-axis shows the temperature, which increases by approximately 12 K per 100 ns. (C) Zoomed in snapshots of model (d) at various simulation times with black arrows pointing to hydrophobic nucleobase-alkyl interactions. The topmost snapshot shows water molecules within 0.25 nm of the residues participating in these hydrophobic interactions that do not obscure the view as light blue spheres.

In the simulations, temperatures that are higher than what is experimentally reasonable are used to promote melting. The reason is that experimental *T*_*m*_ measurements and equilibration take minutes and are performed on bulk solutions with a large number of molecules, whereas the simulations are limited to individual molecules and nanosecond to microsecond time scales. Therefore, temperatures that are much higher than the actual *T*_*m*_ are used to see formation of defects and melting.

Simulation snapshots are shown in Figure 7A and C. Analysis of hydrogen bonding between complementary base pairs is shown in Figure 7B. Supplementary Movie 3 compares trajectory snippets of models (a) and (d).

The heating simulations show that the alkyl-modified duplexes melt more readily than the unmodified one, consistent with our experimental *T*_*m*_ data. This can be explained by alkyl-modified duplexes being more prone to distortions due to hydrophobic interactions the alkyl groups have with themselves and the nucleobases (Figure 7C). Furthermore, comparing models and (d), the heating simulations reveal that the duplex is less structurally and thermally stable when the count of alkyl modifications is higher.

Although molecular dynamics simulations are an extremely useful addition to experiments, this study is also a cautionary tale about their limitations. First, the coarse-grained MARTINI model in ref ^[37]^ which was the original starting point of the current study, cannot predict the stability of highly alkylated DNA structures. Next, single-stranded DNA appears overly stiff in the all-atom AMBER model, while CHARMM produces unrealistic base pair opening in double-stranded DNA. The models need to be chosen carefully for the respective question and predictions need to be experimentally validated.

## Conclusion

In this study, we explored how S-alkyl-phosphorothioates influence the structural and thermal stability of both single-stranded oligonucleotides and double-stranded duplexes. While PS modifications themselves slightly lower the melting temperature *T*_*m*_, the decrease is more drastic after S-alkylation. Both experiments and atomistic MD simulations show qualitatively the same trends, and the simulations reveal two mechanisms that jointly explain the decreasing *T*_*m*_. First, alkylated single-stranded oligonucleotides compact to sequester their hydrophobic groups away from water by the same mechanism that buries hydrophobic amino acid side groups in folded proteins. Moreover, they are less dynamic than unmodified oligonucleotides, further reducing their ability to initiate complex formation with their counter strands. Second, double-stranded DNA with an S-alkylated backbone is structurally and thermally destabilized compared to standard DNA.

Based on our experiences, we offer the following design guidelines:

1. sequence design should not exceed 2-3 S-alkylated PS per helical turn if alkyls are to be placed throughout the entire length of the strand.
2. full helical turn or more of standard nucleotides that are uninterrupted by S-alkyls will improve oligonucleotide hybridization and could allow for incorporating a segment with more than 2-3 S-alkylated PS, if needed.
3. Segments containing S-alkyl PS should be expected to be more flexible due to increased local defects and melting.

While this report shows the limitations of S-Alkyl PS modifications for DNA nanostructures, and our finding that the high degree of functionalization that was suggested in earlier simulations ^[37]^ is unfeasible, they remain highly useful ^[10,14]^ due to their tunability, low cost and ease of functionalization, if the limitations are considered in the structural design.

## Materials and Methods

### Experimental Materials

All phosphorothioate modified DNA oligonucleotides were purchased from Integrated DNA technologies in the desalted form and were suspended in deionized water to form a 1 mM solution. They were then purified in-house with reverse phase-high performance liquid chromatography (RP-HPLC) from Agilent using a Restek Viva 5 µm C4 column (cat. no. 9512525). The oligonucleotides were then dried using a vacuum centrifuge (Eppendorf) followed by resuspension in water to form a 1 mM solution. The yield of the purified oligonucleotides varied between 50-70%. Ethyl (cat. no. I7780), butyl (cat. no. 167304) and decyl iodide (cat. no. 238252) were purchased from Sigma Aldrich, pentyl iodide from Thermo Scientific (cat. no. B20781.22) and heptyl iodide was obtained from TCI chemicals (cat. no. I0320). DMF was obtained from Alfa Aesar (cat. no. 043465).

20% denaturing polyacrylamide gels were hand cast using 8 M Urea (VWR, cat. no. 0568), 10X TBE prepared in-house, 40% acrylamide-bisacrylamide solution (19:1) from Thermo Scientific (cat. no. J60909.K2), 10% ammonium persulphate from VWR (cat. no. M133) and TEMED from Bio Rad (cat. no. 161-0801). GeneRuler Ultra Low Range DNA ladder was purchased from Thermo Fisher Scientific (cat. no. SM1211). 5% native gels with 5 mM MgCl_2_ were hand cast using 10X TBE, 40% acrylamide-bisacrylamide solution (29:1), 10% APS, TEMED and MgCl_2_ hexahydrate (VWR, cat. no. BDH9244). All gels were run in 1X TBE (100 mM TRIS, 100 mM Boric Acid, 2 mM EDTA, pH 8.3). Tris(hydroxymethyl)amino methane was purchased from Sigma Aldrich (cat. no. 252859), Boric acid (cat. no. BDH9222) and disodium salt of EDTA were purchased from VWR (cat. no. BDH4616). SYBR gold (cat. no. S11494) and SYBR green (cat. no. S7563) DNA gel stains were purchased from Invitrogen by Thermo Fisher Scientific. HEPES (cat. no. J848) was purchased from VWR, MgSO_4_.7H_2_O (cat. no. M2773) and Na_2_SO_4_ (cat. no. 71959) were purchased from Sigma Aldrich. 6X native DNA loading dye (cat. no. B7024S) and 100 bp DNA ladder (cat. no. N3231) were purchased from New England Biolabs. Zymo-Spin IICR columns (cat. no. C1078), Oligo binding buffer (cat. no. D4060-1-40) and DNA wash buffer (cat. no. D4003-2-48) were purchased from Zymo research.

### Experimental Methods

#### S-Alkylation of PS DNA

A solution of HPLC purified PS oligonucleotides (5 µL, 1 mM) was mixed with Tris HCl pH 8.0 (2 µL, 30 mM) in a DNA-low bind microcentrifuge tube. This step maintains the deprotonated state of the PS group making it suitable for nucleophilic attack at the α-carbon of the alkyl iodide. To this solution, anhydrous, amine-free DMF (18 µL) was added and mixed well. Care must be taken to ensure that the DMF is amine-free, else the dimethyl amine contaminations will interfere in the nucleophilic substitution causing undesirable side products and fragmentation (Figure S10, Supporting Information**Error! Reference source not found**.). Next, alkyl iodide (5 µL) was added to make a final volume of 30 µL and incubated in a thermomixer at 65 °C, 1000 rpm for 4 h.

Post-reaction, the solvent was evaporated in a vacuum centrifuge, and the pellet was redissolved with EDTA solution (50 µL, 0.1 M, pH 8.0), heated to 90 °C for 5 min. Hot EDTA solution was found to be necessary to efficiently re-dissolve the DNA pellet. Solubility in organic solvents was not tested because lipids could also be solubilized in organic solvents thereby hindering subsequent nanodisc formation. The alkylated DNA was then separated from the remaining iodo compound by G-25 size exclusion spin columns. Yields of the purified product were measured by UV absorbance and were about 50-60%. Denaturing gel and HPLC analysis show that this method led to quantitative conversion of the starting PS to its S-alkylated form and hence the product was not further purified. Mass spectrometry was done in refs. ^[10,14,39]^ which confirmed that N-alkylation was not observed, and exclusive S-alkylated products were obtained.

Refer to Table 1 for the sequences of the PS oligonucleotides.

**Table 1.**
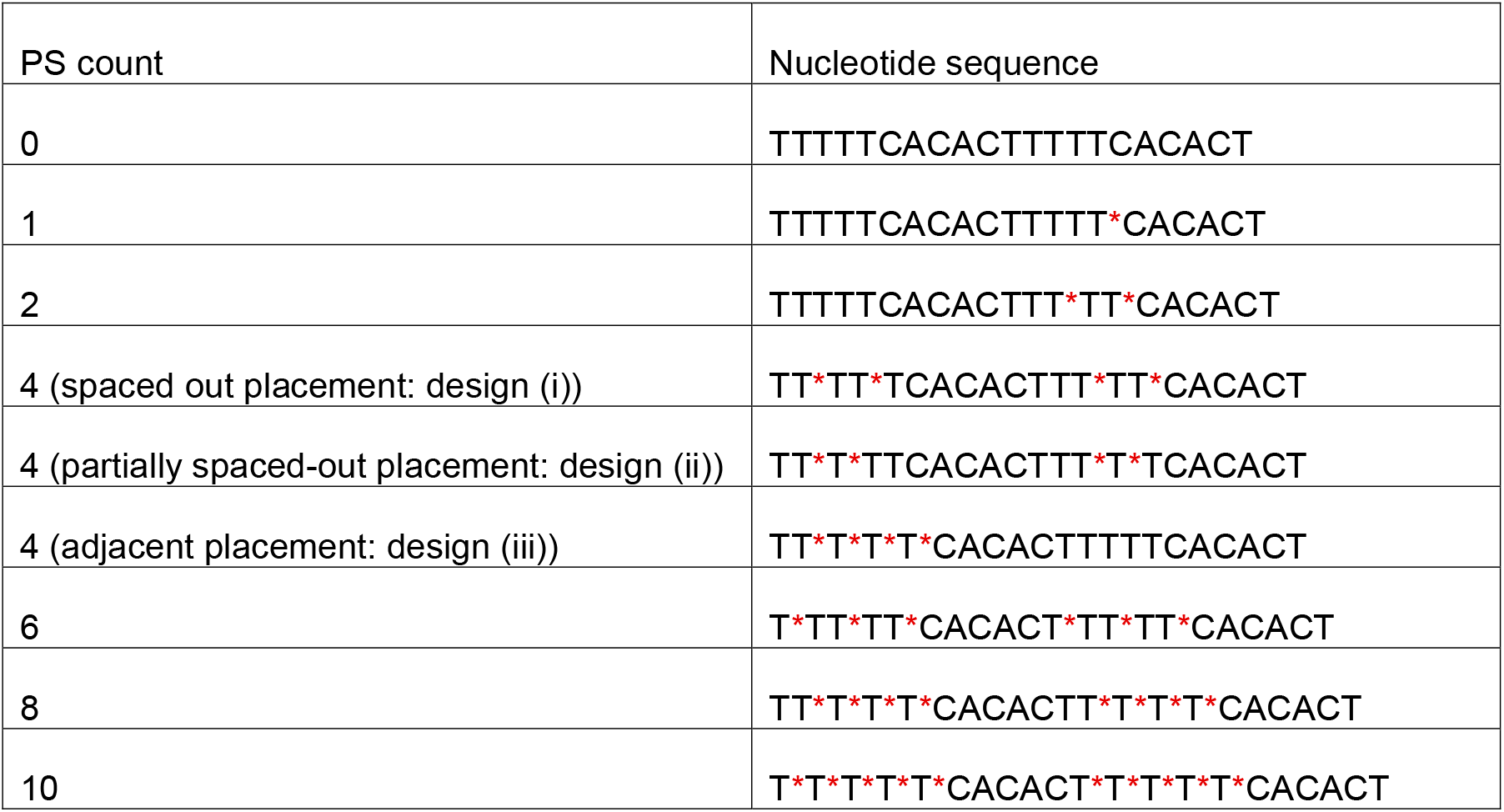
Nucleotide sequences of the PS oligonucleotides. The asterisks (*) indicate the placement of PS groups.

#### Denaturing polyacrylamide gel electrophoresis

20% denaturing urea polyacrylamide gels were cast using 40% acrylamide-bisacrylamide (5 mL), 10X TBE (1 mL), urea (4.8 g), 10% APS (100 µL), TEMED (10 µL) and water to make up the total volume to 10 mL. Gels were polymerized for 30-60 min before samples were loaded. The starting PS (7.5 µL, 20 pmol) and the alkylated product (7.5 µL, 20 pmol) were each mixed with 4X denaturing loading dye (2.5 µL, containing 100% formamide, 10 mM NaOH, 0.01% bromophenol blue, 0.02% xylene cyanol) and loaded in the sample wells. The GeneRuler ultralow range DNA ladder (1 µL, 0.01 µg/mL) was also mixed with the denaturing loading dye and applied to sample wells as a reference.

Electrophoresis was performed at 200 V for 65 min at 65 °C. Denaturing gels are run at high temperatures and in the presence of urea to preserve the DNA in the denatured condition without formation of secondary structures.

#### Formation of single-stranded circular scaffold and double-stranded minicircles

Single-stranded circles were synthesized according to a previously published protocol.^[14]^ Briefly, 147-nt linear oligonucleotides were phosphorylated at their 5’ ends using T4 phosphonucleotide kinase; their respective 5’ and 3’ ends were then splint ligated using T4 DNA ligase. The sequence of the single-stranded 147-mer DNA is: /5Phos/(TGTGAAAAAAGTGTGAAAAAG)_7_.

Uncircularized strands and the splints were removed by digestion with Exonuclease I and III. Pure circular product was obtained after affinity column purification (DNA clean, Zymo Research) which separates the pure circular ssDNA scaffold from enzymes and nucleotides, followed by yield estimation using absorbance measurements.

Double-stranded (ds) minicircles were formed by annealing the circular scaffold (500 pmol) with the complementary unmodified or alkylated oligonucleotides (12.5 nmol) in 5 mM MgCl_2_ by heating to 80 °C and slowly cooling them to 25 °C over an hour. The ds-circles were purified with 50 kDa molecular weight cut-off (MWCO) filters in a buffer containing 50 mM HEPES, 5 mM MgSO_4_ and 100 mM Na_2_SO_4_. The sample was washed 5-7 times with buffer (500 µL) and spun at 10000 rcf for 5 min. Finally, the filter was inverted and the sample was retrieved by spinning at 10000 rcf for 3 min. Yields were determined by measuring absorbance at 260 nm and the ds circles were analyzed by native polyacrylamide gel electrophoresis (PAGE) or AFM.

#### Native polyacrylamide gel electrophoresis

5% native polyacrylamide gels were cast using 40% acrylamide-bisacrylamide (1.25 mL), 10X TBE (1 mL), 200 mM MgCl_2_ (250 µL), 10% APS (100 µL), TEMED (10 µL) and water (7.39 mL) to make up the total volume to 10 mL. Gels were allowed to polymerize for 30-60 min before samples were loaded. The single-stranded circular scaffold (10 µL, 0.5 pmol) was mixed with the commercial 6X native loading dye (2 µL, containing 2.5% Ficoll-400, 10 mM EDTA, 3.3 mM Tris-HCl, 0.08% SDS, 0.02% Dye 1, 0.001% Dye 2, pH 8.0 at 25 °C) was loaded in the sample wells. Since SYBR gold stains dsDNA better, 0.25 pmol of ds-rings were loaded into gels. The 100 bp DNA ladder (1 µL, 50 µg/mL) was also mixed with the native loading dye and applied to sample wells as a reference. Electrophoresis was performed at 100 V for 65 min at 4 °C.

#### Denaturing and native PAGE staining and imaging

When electrophoresis was complete, the gel was stained in 1X SYBR gold in 1X TBE with 5% ethanol to prevent the stain from sticking to the plastic staining tray. Gels were then imaged on a GE Typhoon FLE 9500 gel scanner using a 473 nm excitation laser and 510 nm long pass emission filter. The photomultiplier tube gain was set to 500 and the pixel size was 50 µm. Images were analyzed by the Fiji ImageJ image analysis software.

#### Atomic force microscopy

Polyornithine (20 µL, 0.01%, Sigma Aldrich, cat. no. P3655) solution was added onto freshly cleaved mica and incubated for 2 min. The mica surface was then washed with Milli-Q water (∼10 mL) and dried with a stream of N_2_ gas. Unmodified and alkylated ds-rings (10 µL, 20 nM) were deposited onto the mica (Ted Pella, 9.9 mm diameter) and incubated for 3–5 min. The solution was wicked away and the surface was washed twice with Milli-Q water (1 mL) after which water (60 µL) was added on to the sample surface. Water (20 µL) was also added onto the ScanAsyst Fluid+ tip (Bruker, cat. no. p-3728) and imaging was performed in liquid using the peak force tapping mode in a highspeed atomic force microscope (Bruker, Dimension Fastscan bio). Images were analyzed using Nanoscope analysis software.

#### *T*_*m*_ analysis

A 21-mer DNA which is the exact complement of the alkylated oligonucleotide was designed (sequence: AGTGTGAAAAAGTGTGAAAAA). The 21-mer DNA (1 µL, 200 µM) was mixed with a 5-fold molar excess of the complementary unmodified or alkylated oligonucleotide (1 µL, 1 mM) in 1X TE buffer with 5 mM MgCl_2_. To this 10X SYBR green (3 µL, such that final concentration was 1X) was added to record real-time change in fluorescence upon DNA hybridization. The total reaction volume was 30 µL. Additionally, samples were also layered with oil to prevent evaporation. Samples were placed in a real-time magnetic induction cycler (Biomolecular Systems) and heated to 85 °C for 2 min through magnetic induction followed by cooling to 40 °C over an hour using fan-forced air. Fluorescence from the DNA-binding dye changed as the temperature decreased due to formation of dsDNA and was recorded with high-sensitivity photodiodes for each dedicated excitation/emission channel. micPCR software was used to calculate the first derivative of change in fluorescence with change in temperature to obtain the *T*_*m*_ of the duplex.

### Atomistic Molecular Dynamics Simulations

#### Parameterization of non-standard nucleotides

Force field parameters for modified nucleic acids, such as those with phosphorothioates, are still under development.^[50,51]^ Due to the lack of reliable parameters for S-alkyl-phosphorothioates, the modified nucleotides were instead constructed with O-alkyl-phosphotriesters like in previous studies.^[39,52]^ Calculating and validating reliable parameter sets for S-alkyl-phosphorothioates from *ab initio* quantum mechanics calculations is beyond the scope of this paper. Considering the large overall size of these functional groups, the relative difference between S-alkyl-phosphorothioates and O-alkyl-phosphotriesters is negligible. Moreover, the charge elimination and the additional hydrophobic surface after alkylation can be expected to dominate the behavior of the molecule in the simulations. Lastly, the oxygen and sulfur atom in the DNA backbone have the same valency, similar bond angles, and differ in size and electronegativity only slightly in a functional group containing 12-36 atoms. Thus, this parameterization choice seems acceptable.

For both the AMBER Parmbsc1 force field^[46]^ and the CHARMM36 force field,^[53]^ the modified central nucleotide fragments were constructed by using two elementary building blocks: dimethyl ethyl- or decyl-phosphate and a deoxynucleoside.

For the Parmbsc1 force field, the modified nucleotide fragments were parametrized with the Parmbsc1 force field with the CUFIX set of corrections for nonbonding interactions.^[44]^ Restrained electrostatic potential (RESP) charges^[54]^ (HF/6-31G*//HF/6-31G*) for the modified nucleotide fragments were obtained with the PyRED program (version January 2025) available in R.E.D. Server^[55]^ using Gaussian 16 (Rev. C.01).^[56]^

For the CHARMM36 force field, the force field parameters for the charge-neutral nucleotide variants were obtained using the CGenFF server.^[57]^ All the penalty values reported by the CGenFF server were less than 10, indicating the parameterization is fair. An additional charge of +0.013 was distributed among the partial charges of the 5’ and 3’ carbon and oxygen atoms, i.e. the connecting atoms of the two elementary building blocks, so that the total charge of the nucleotide is zero. The atom types of the modified nucleotide fragments, except for at the alkyl-phosphate group, were changed to NA36 atom types, matching the atom types of their analogous standard nucleotide residues (Figure S11, Supporting Information). The missing bond, angle, and dihedral parameters caused by the linkage of CGenFF36 and NA36 atom types were obtained by analogy. Gromologist^[58]^ was used as a tool to find analogous parameters.

#### MD protocols

All MD simulations were performed in GROMACS (version 2024.3)^[59]^ on GPUs at the Ohio Supercomputer Center.^[60]^ CUFIX corrections were applied to both the Parmbsc1 and CHARMM36 force field. CUFIX corrections were applied to the CHARMM36 force field for GROMACS using charmm2gmx.^[61]^

The TIP3P model^[62]^ was used to represent water in Parmbsc1 simulations, and the CHARMM-modified TIP3P model in CHARMM36 simulations. The DNA was centered in a cubic box and solvated with explicit water molecules. The DNA was placed at least 1 (resp. 1.2) nm from the edge of the box for Parmbsc1 (resp. CHARMM36) simulations. All systems were neutralized with sodium ions, followed by the addition of NaCl salts to a concentration of 150 mM.

All minimizations and simulations were performed using periodic boundary conditions with the particle mesh Ewald^[63]^ method used for treating long-range electrostatics. The bonds containing hydrogen atoms were constrained using the LINCS algorithm,^[64]^ and a 2 fs time step was used for all MD simulations. All minimizations were performed with the steepest-descent algorithm until the maximum force was less than 1000 kJ/mol/nm.

After minimization, a 100 ps NVT equilibration run was performed at 300 (resp. 298) K for Parmbsc1 (resp. CHARMM36) simulations. This was followed by a 100 ps NPT equilibration run at 1 bar pressure. Both equilibration runs involved harmonic restraints of 1000 kJ/mol/nm^2^ on the heavy, i.e. non-hydrogen, atoms of the DNA. Short range non-bonded interactions were cut off at 1 (resp. 1.2) nm for Parmbsc1 (resp. CHARMM36) simulations.

Constant temperature production runs were performed at a constant pressure of 1 bar and a temperature of 300 (resp. 298) K for Parmbsc1 (resp. CHARMM36) simulations using a V-rescale thermostat^[65]^ and a C-rescale barostat.^[66]^ The duration of the production runs were 600 ns.

Production runs with gradual heating used the “simulated annealing” option available in GROMACS and the Parmbsc1 force field. The temperature of the system was increased from 300 K to 440 K within 1200 ns. The initial configurations were the final configurations of the corresponding constant temperature production runs. A V-rescale thermostat and a C-rescale barostat were applied.

#### Construction of DNA structures

A PDB file defining the atomic model of 21 base pair double-stranded DNA was generated using the BIOVIA Discovery Studio 2024^[67]^ software according to the experimental nucleotide sequence. From this model, a single-stranded DNA model was constructed by removing the complementary strand.

From these two unmodified models, eight alkyl-modified DNA models were constructed, i.e. four for both single- and double-stranded DNA, using the tLEaP Ambertools^[68]^ module. That is, tLEaP was used to add the missing alkyl groups to the modified models according to the force field libraries generated by PyRED. The modifications were placed on alternating non-bridging oxygens because experimentally there is no control over which non-bridging oxygen atom is replaced with a sulfur atom.

All DNA models were saved as a PDB format file using tLEaP. Additionally, all the alkyl-modified DNA models were saved as an Amber topology and coordinate file using tLEaP. GROMACS topologies and coordinate files were generated from these Amber ones using acpype (v. 2023.10.27);^[69]^ these GROMACS files were used as initial structures for only the Parmbsc1-based simulations of alkyl-modified DNA.

For CHARMM36-based simulations, since the PDB files of the DNA models outputted by tLEaP have AMBER naming conventions, atom names and terminal residue names were changed, using a text editor, to comply with the CHARMM36 force field. The GROMACS topologies were then created using the GROMACS program `pdb2gmx`.

#### Analysis

The radius of gyration for each snapshot was measured using the GROMACS program ‘gyrate’. The twist at each base pair step was calculated using the softwares 3DNA ^[70]^ and do_x3dna.^[71]^ The twist averaged over the base pairs, excluding the three terminal ones at each end of the duplex, was calculated using a Python script written in-house. All time evolutions of average twist and gyration radius were smoothed using a double moving average: initially with a 25-ns window, then with a 5-ns window.

The geometrical criterion defined in the GROMACS manual ^[72]^ is used to determine the existence of a hydrogen bond. That is, the donor-acceptor distance must be less than or equal to 0.35 nm and the hydrogen-donor-acceptor angle must be less than or equal to 30°. Only one donor-hydrogen-acceptor triplet, i.e. the one where the acceptor is a nitrogen atom, of a Watson-Crick base pair is analyzed to determine if it is intact or not. The donor-acceptor distance and hydrogen-donor-acceptor angle were measured for each snapshot using the GROMACS utilities ‘distance’ and ‘angle’, respectively.

Python scripts were written in-house to plot analysis data. Trajectories were visualized by means of the PyMOL Molecular Graphics System (v3.1.4.1).^[73]^

## Data and materials availability

Raw experimental or simulation data is available upon request.

## Author Contributions

‡S.C. and R. B. contributed equally. T.L.S., S.C. and R.B. conceived the study; S.C. performed experiments; R.B. performed all-atom MD simulations; F.F. created the MD protocols; S.C., R.B. and T.L.S. analyzed and visualized the data; T.L.S. acquired funding and supervised the study; S.C., R.B. and T.L.S. wrote the initial draft; all authors revised and expanded the manuscript.

## Notes

The authors declare no competing financial interest.

## Acknowledgments

We thank Dr. Aleksei Aksimentiev, Dr. Christopher Maffeo, Dr. Jejoong Yoo, Dr. Hemani Chhabra, and Dr. François-Yves Dupradeau for helpful discussions.

## Table of Contents Graphic

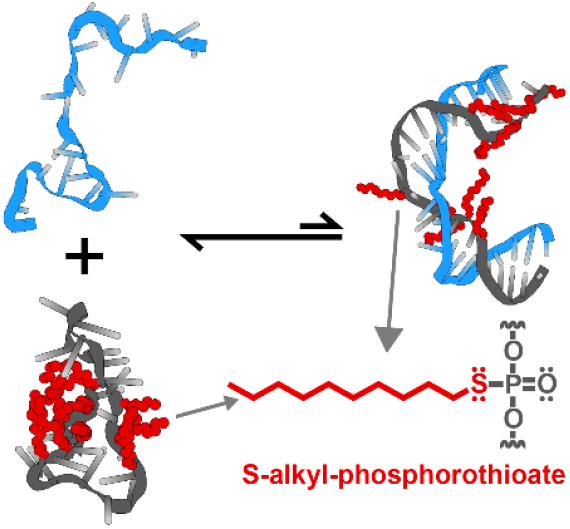

## Table of Contents

S-Alkylation of phosphorothioates enhances hydrophobicity of DNA and enables interactions with lipids but destabilizes dsDNA. Based on experiments and molecular dynamics simulations, two mechanisms are proposed for destabilization: compaction of ss oligonucleotides and melting of dsDNA more easily due to increased hydrophobicity. This paper gives practical guidelines for the S-alkylation of phosphorothioate DNA in future structural DNA nanotechnology experiments

