## Supporting information for "S-Alkyl-Phosphorothioate Modifications Reduce Thermal and Structural Stability of DNA Duplexes"

‡Contributed equally

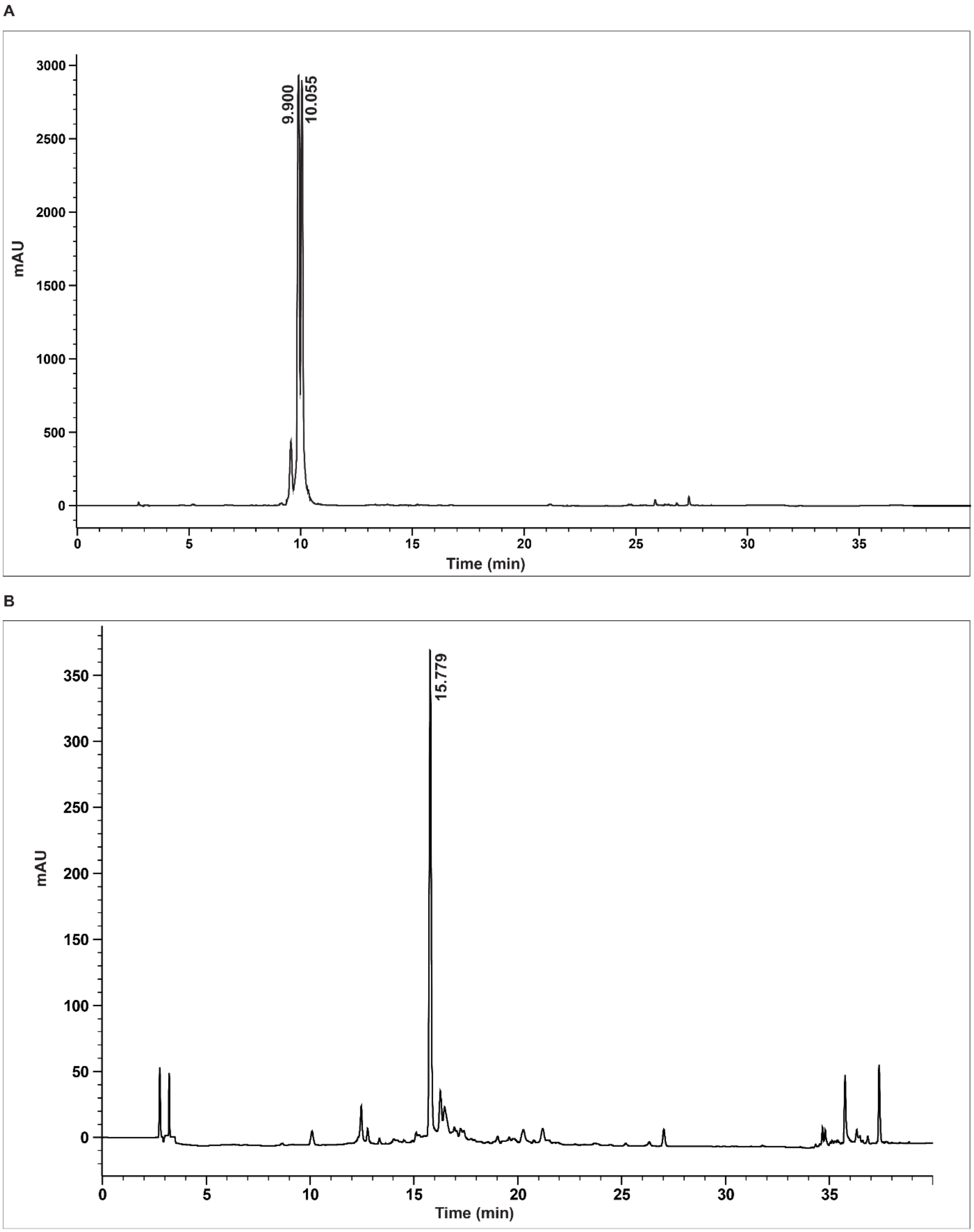


**Figure S1**. HPLC chromatograms of alkylated oligonucleotides. (A) HPLC chromatogram of starting 1 PS oligonucleotide with retention time at ~10 min. DNA appears as two peaks, or two different diastereomers, from the oxidation step with the sulfurizing Beaucage reagent.[1] (B) 1 PS DNA coupled with decyl iodide with a shift in retention time to ~15 min due to increased hydrophobicity from the alkyl group. The absorbance is different in the two chromatograms because the sample amounts loaded were different.

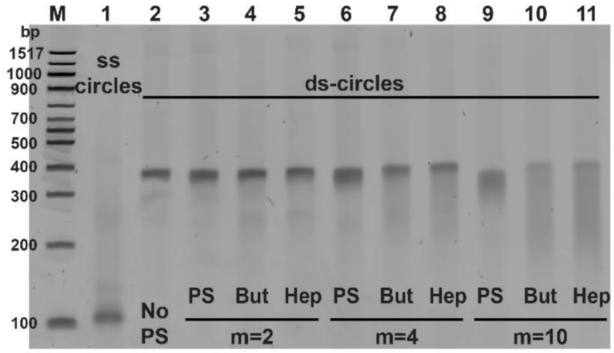


**Figure S2.** Native electropherogram of ds-minicircles with different number and length of alkyl chains. For this, a 147 nt scaffold was hybridized with 7 identical complementary strands with varying number of PS groups (m) and length of alkyls. The excess of short oligonucleotides runs out of the gel. M: Marker (100 bp dsDNA ladder); Lane 1: ss-scaffold; Lane 2: dscircle with no PS groups; Lane 3: ds circle with 2PS groups, not alkylated; Lane 4: dscircle with 2PS groups, butyl modified; Lane 5: dscircle with 2PS groups, heptyl modified; Lane 6: dscircle with 4PS groups, not alkylated; Lane 7: ds circle with 4PS groups, butyl modified; Lane 8: dscircle with 2PS groups, heptyl modified; Lane 9: dscircle with 10PS groups, not alkylated; Lane 10: ds circle with 10PS groups, butyl modified; Lane 11: ds circle with 10PS groups, heptyl modified.

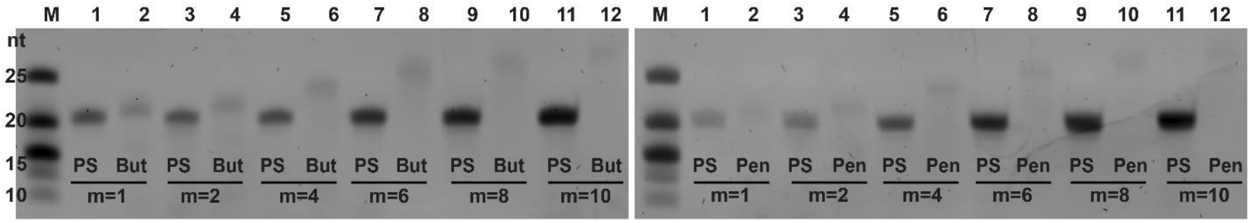


**Figure S3**. Denaturing gel electropherogram of alkylated oligonucleotides. Left gel: with m=1-10 PS, before and after reaction with butyl iodide (n=3). Right gel: with m=1-10 PS, before and after reaction with pentyl iodide (n=4). Odd numbered lanes have starting PS DNA, and even numbered lanes have the corresponding alkylated product. M: Ultra-low range DNA ladder.

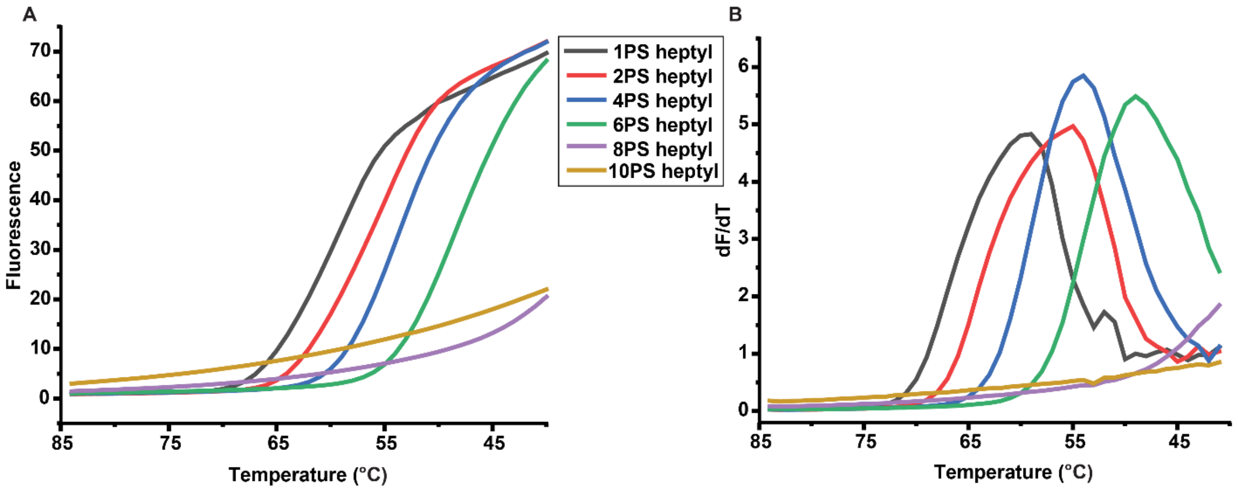


**Figure S4.** *$T_m$* analysis for alkylated DNA. (A) Raw data from qPCR cycler measuring the real-time change in fluorescence (Y-axis) with change in temperature from 85 °C to 40 °C indicated in the X-axis (B) Melting temperature (*$T_m$*) obtaining by plotting first derivative of change in fluorescence with respect to temperature against change in temperature for all alkylated DNA with m=1-10 and n=6 (heptyl).

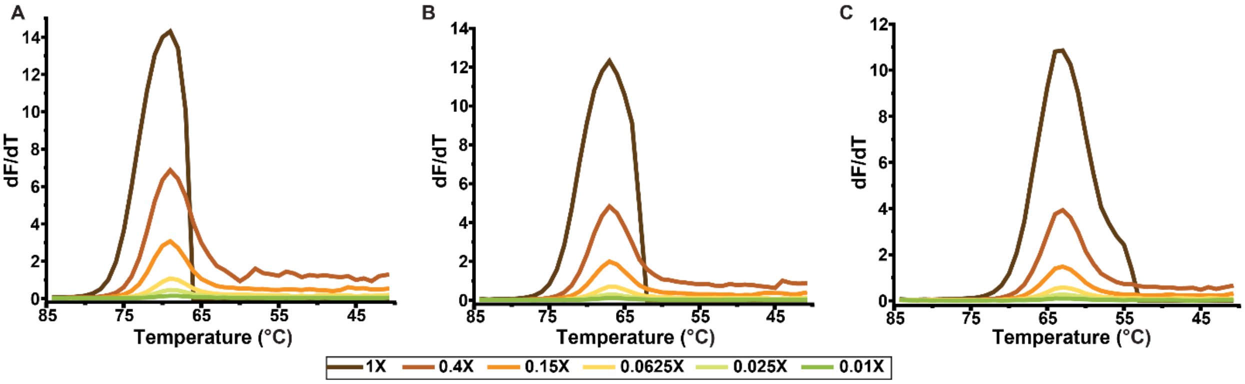


**Figure S5.** Influence of SYBR green concentrations from 0.01X to 1X on $T_m$ of different PS DNA. (A) $T_m$ of no PS DNA at 69 °C. (B) $T_m$ of 4PS DNA at 66 °C. (C) $T_m$ of 10PS DNA at 62 °C (X-axis - change in temperature from 85 °C to 40 °C)

| Measured Tm (°C) | Alkylated PS DNA | Number of modifications | DNA with mismatches | Predicted Tm (°C) |
|---|---|---|---|---|
| 69.0 | 5'-TTTTTCACACTTTTTCACACT-3'<br>3'-AAAAAGTGTGAAAAAGTGTGA-5' | No mismatch/ alkylation | 5'-TTTTTCACACTTTTTCACACT-3'<br>3'-AAAAAGTGTGAAAAAGTGTGA-5' | 67.6 |
| 61.0 | 5'-TTTTTCACACTTTTT$^H$CACACT-3'<br>3'-AAAAAGTGTGAAAAAGTGTGA-5' | 1 T-C mismatch/ 1-heptyl | 5'-TTTTTCACACTTTTCCACACT-3'<br>3'-AAAAAGTGTGAAAAAGTGTGA-5' | 62.5 |
| 58.0 | 5'-TTTTTCACACTTT$^H$TT$^H$CACACT-3'<br>3'-AAAAAGTGTGAAAAAGTGTGA-5' | 2 T-C mismatches/ 2-heptyl groups | 5'-TTTTTCACACTTCTCCACACT-3'<br>3'-AAAAAGTGTGAAAAAGTGTGA-5' | 56.8 |
| 51.5 | 5'-TT$^H$TT$^H$TCACACTTT$^H$TT$^H$CACACT-3'<br>3'-AAAAAGTGTGAAAAAGTGTGA-5' | 4 T-C mismatches/ 4-heptyl groups | 5'-TCTCTCACACTTCTCCACACT-3'<br>3'-AAAAAGTGTGAAAAAGTGTGA-5' | 50.1 |
| 54.0 | 5'-TT$^H$T$^H$TTCACACTTT$^H$T$^H$TCACACT-3'<br>3'-AAAAAGTGTGAAAAAGTGTGA-5' | 4 T-C mismatches/ 4-heptyl groups (two placed together) | 5'-TCCTTCACACTTCCTCACACT-3'<br>3'-AAAAAGTGTGAAAAAGTGTGA-5' | 56.0 |
| 57.5 | 5'-TT$^H$T$^H$T$^H$T$^H$CACACTTTTTCACACT-3'<br>3'-AAAAAGTGTGAAAAAGTGTGA-5' | 4 T-C mismatches/ 4-heptyl groups (all four placed together) | 5'-TCCCCCACACTTTTTCACACT-3'<br>3'-AAAAAGTGTGAAAAAGTGTGA-5' | 64.4 |

**Figure S6.** Comparison of measured $T_m$ of alkylated and $T_m$ of mismatched DNA. Column 3 indicates the number and kind of modification. Columns 2 and 4 indicate 21mer DNA with identical sequences and alkylation/mismatch at the same positions. A "T to C" mismatch was done here as that is the most discriminating mismatch. The $T_m$ of DNA with mismatches were analyzed at 5 mM $Mg^{2+}$ and 10 mM $Na^+$ concentrations *mfold*.[2]

Qualitatively, the trends are similar: DNA without mismatches or alkylations showed the highest $T_m$ as predicted. Increasing number of mismatches/alkyl groups decreased $T_m$ progressively. DNA with mismatches spaced out (Row 5) shows a lower $T_m$ than DNA with mismatches placed together (Row 7) similar to the pattern observed in alkylated DNA with identical PS organization (Figure 4B). This indicates that placing all modifications together on one end of the helix leaves the rest of the DNA undisturbed, leading to a higher $T_m$ whereas spacing them out causes frequent distortions in the helical structure and the cooperative binding of neighboring base pairs, thereby lowering the $T_m$. Moreover, spaced out alkyls also compacted the oligonucleotide more (Figure 5).

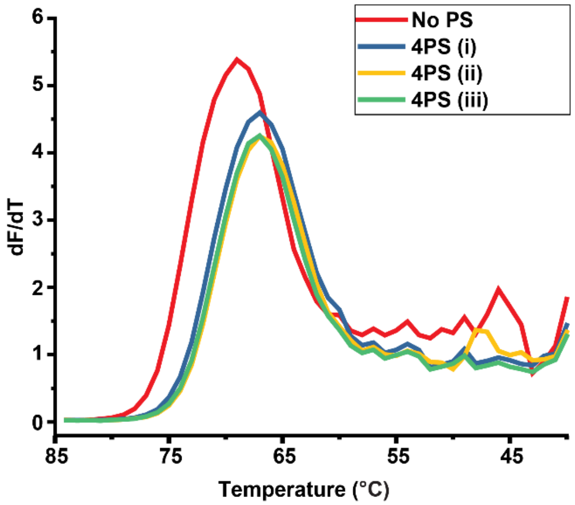


**Figure S7.** $T_m$ of DNA containing same number of PS groups at different positions in the strand. Red: DNA with no PS group with $T_m$ at 69 °C; Blue, yellow, green; DNA with different positions of PS groups (refer to Figure 4A for design) showing identical $T_m$ at 66 °C (X-axis - change in temperature from 85 °C to 40 °C).

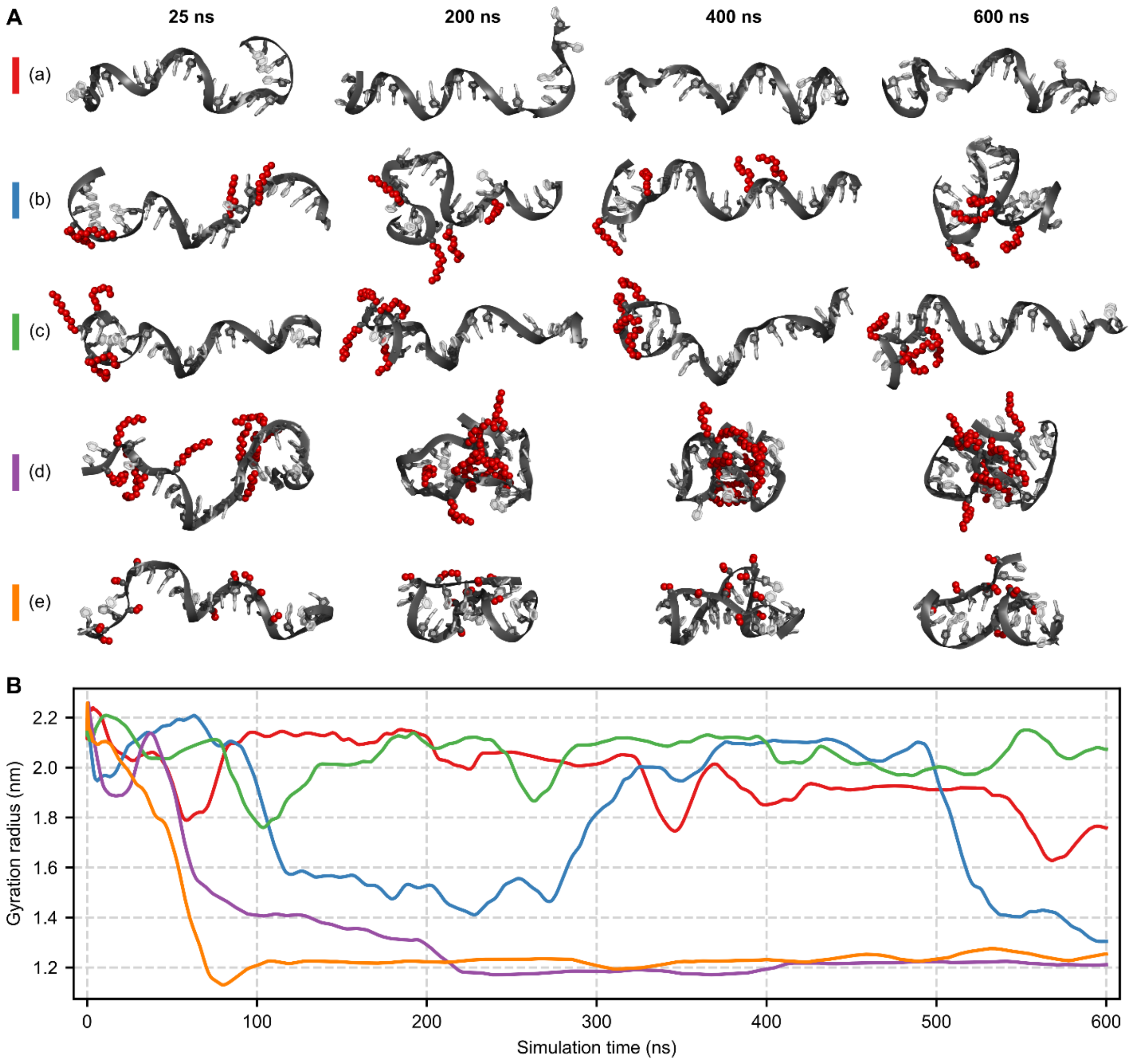


**Figure S8.** MD simulations of unmodified and alkyl-modified single-stranded DNA (ssDNA) using the Parmbsc1 force field. (A) Simulation snapshots at different times of the trajectory. (B) Radius of gyration of each model plotted as a function of simulation time. The color code is shown in panel (A).

Among all the alkyl-modified models, model (c) behaves the most like the unmodified model (a). This can be explained by model (c) having the longest stretch of unmodified nucleotides among the alkyl-modified models. Moreover, ssDNA compacts more when the modifications are placed spaced-out, as seen in the comparison between models (b) and (c). Lastly, a higher count of modifications also causes ssDNA to compact more and be less dynamic. That is, the extensively alkyl-modified models (d) and (e) are the most compact and least dynamic models.

Even though the Parmbsc1 force field represents unmodified ssDNA as unnaturally expanded, alkylation has a similar effect as in the CHARMM36-based simulations (Figure 5). That is, alkylation promotes compaction in both force fields.

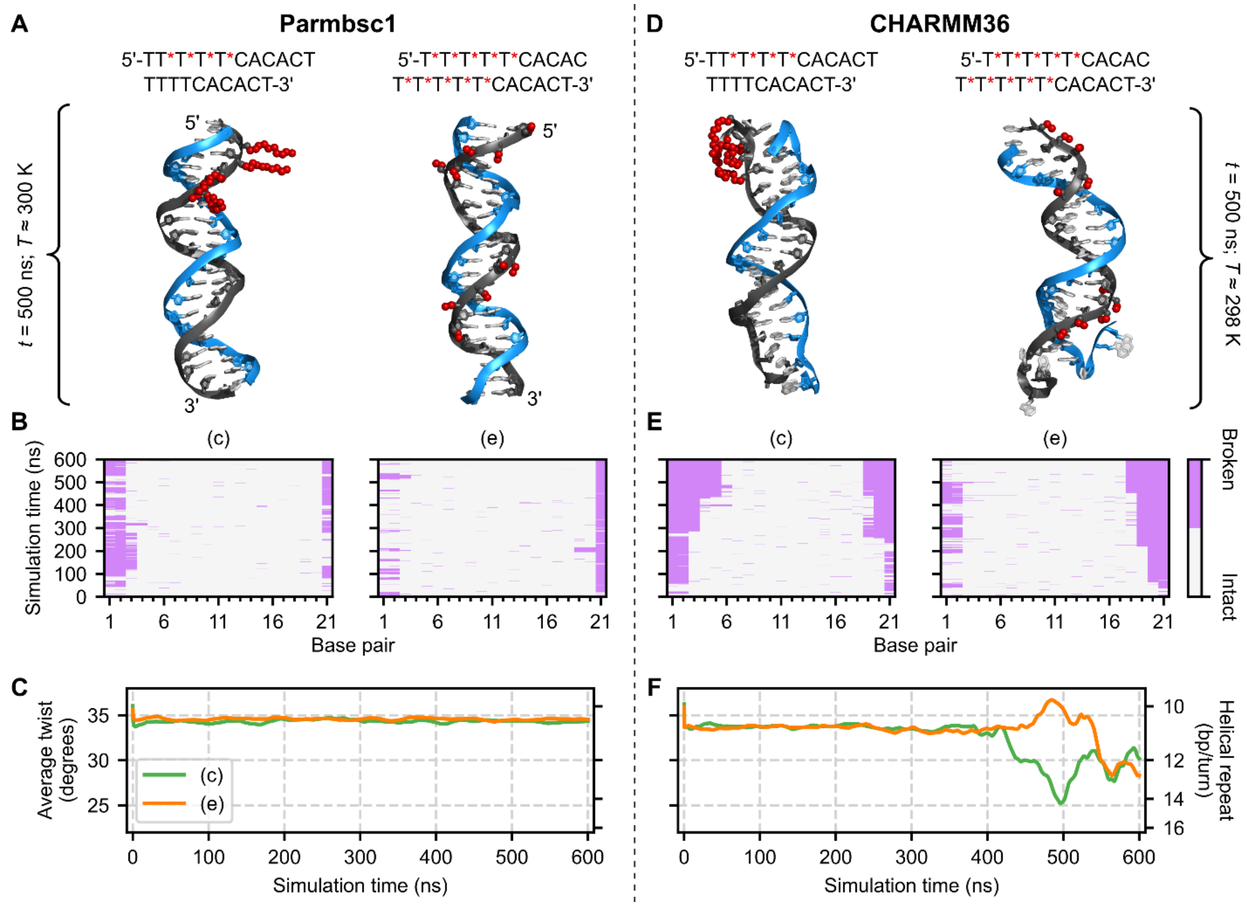


**Figure S9.** Constant temperature MD simulations of alkyl-modified double-stranded DNA. The left column (i.e. A-C) shows Parmbsc1-based simulations, the right (i.e. D-F) CHARMM36-based simulations. (A, D) Simulation snapshots at simulation time *t* and system temperature *T*. (B, E) Analysis of the existence of hydrogen bonding between each Watson-Crick base pair for each frame. (C, F) Twist averaged over the base pairs, excluding the three terminal ones on each end of the duplex, plotted as a function of simulation time.

In combination with the results displayed in Figure 6, both the Parmbsc1- and CHARMM36-based simulations suggest that the stability of the duplex structure depends primarily on the type of modification. That is, the duplexes are less stable the longer the alkyl groups are. For example, comparing the decyl-modified model (d) from Figure 6 and its ethyl counterpart (e), duplexes with longer alkyl groups experience more end-fraying in Parmbsc1-based simulations. Moreover, as seen in the comparison between the same models in CHARMM-based simulations, duplexes with longer alkyl groups experience more backbone distortions.

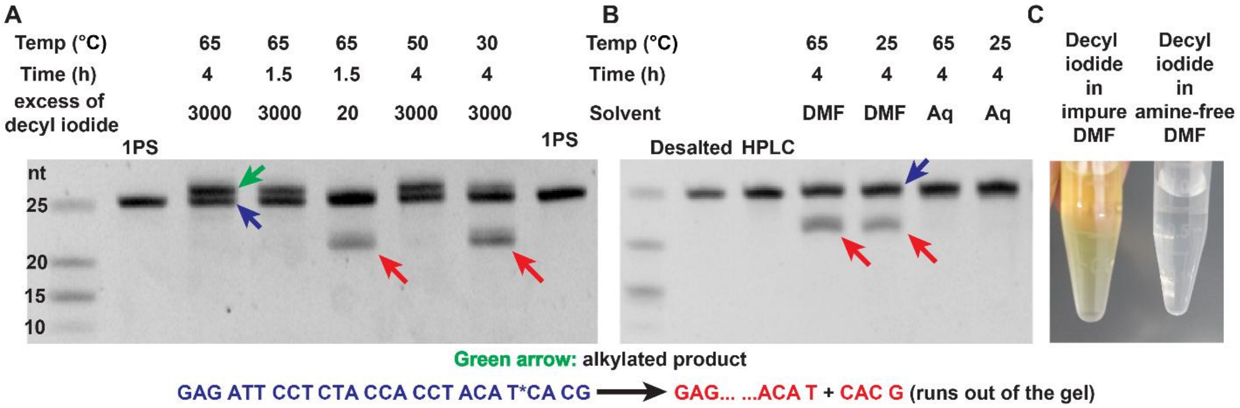


**Figure S10.** Cleavage at the PS is a side reaction. A 26-mer PS oligonucleotide (GAG ATT CCT CTA CCA CCT ACA T*CA CG) is reacted at different conditions with decyl iodide (A) or incubated in different solvents without decyl iodide (B). Depending on the conditions, the 26-mer DNA can fragment at the PS group into a 22-mer and a 4-mer which runs out of the gel. The splitting of the band in (A) around 22 nt likely indicates a fragment with or without the decyl modification. Fragmentation was observed with: low absolute concentration of DNA (in the low µM range); low alkyl iodide excess; temperatures below 50 °C; and poor DMF quality. Fragmentation can be avoided by: running reactions at to 65 °C; high concentrations of DNA; a high excess of alkyl iodide; and using anhydrous, amine-free DMF. Lane 1 in (A) and (B): Ultra-low range DNA ladder. The product conversion in (A) was not quantitative because the alkylation and workup protocol was not yet optimized at that time.

(B) Fragmentation of PS oligonucleotide occurs in DMF also without the addition of alkyl iodide. No significant fragmentation occurs in water, implying that traces of dimethylamine in DMF cause the fragmentation. Cleavage of the PS oligonucleotide in DMF happens irrespective of the quality of the solvent (data not shown) as hydrolysis of DMF also produces dimethylamine. (C) Pure decyl iodide in older DMF with dimethylamine contaminations and in anhydrous, amine-free DMF. The yellow color likely stems from the attack of dimethyl amine on decyl iodide that releases the yellow HI. A high alkyl iodide concentration therefore likely quenches traces of dimethyl amine protecting PS oligonucleotides from cleavage.

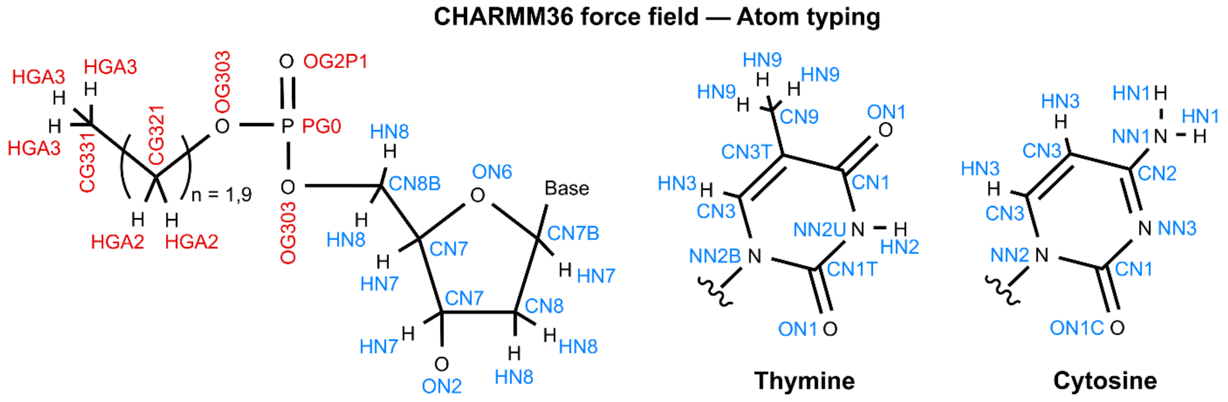


**Figure S11.** Atom typing of the internal modified nucleotide fragments for the CHARMM36 force field. NA36 atom types are shown in blue, CGenFF36 in red.

3. Captions to Supplementary Movies

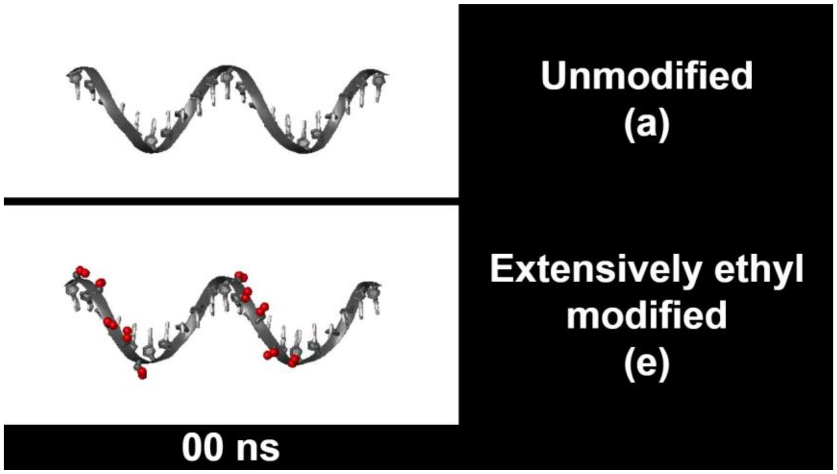


**Supplementary Movie 1.** All-atom MD CHARMM36-based simulations of the unmodified (a) and extensively ethyl modified (e) models from Figure 5A and B. The movie shows the first 17.5 ns of the simulation trajectories. Model (e) quickly collapses to shield its hydrophobic alkyl groups from water.

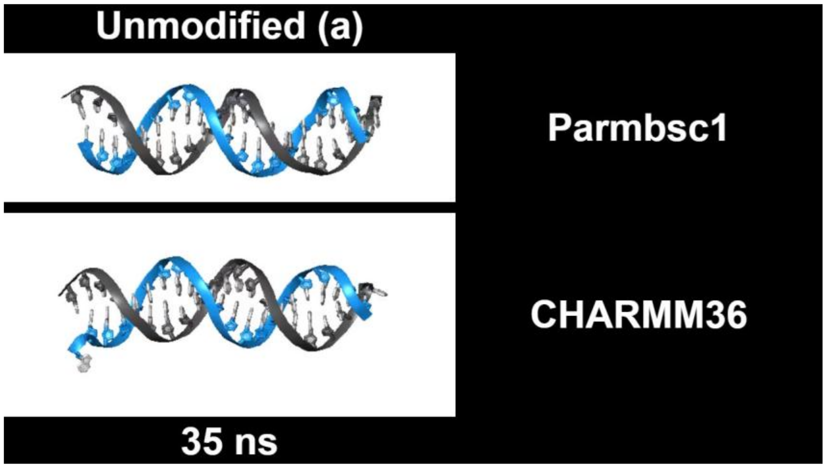


**Supplementary Movie 2.** All-atom MD simulations of the unmodified model (a) from Figure 6 using either the Parmbsc1 or CHARMM36 force field. The movie starts at simulation time 35 ns and ends at 60 ns. The CHARMM36-based simulation begins to show internal base pair opening at approximately the 42 ns time step. Two passes of the PyMOL function `smooth`, i.e. a running average of coordinates, were applied to both trajectories.

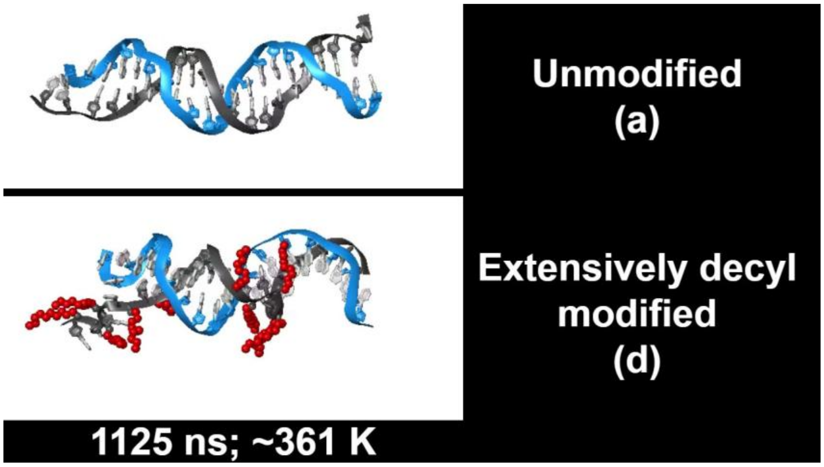


**Supplementary Movie 3.** All-atom MD Parmbsc1-based simulations of the unmodified (a) and extensively decyl modified (d) models from Figure 7 with gradually increasing system temperature. The movie first shows the simulation time window 1125 ns to 1133.5 ns, cuts to black, and then 1725 ns to 1733.5 ns. The unmodified model (a) maintains the duplex structure better than the extensively decyl modified model (d).
